# Inferring Metabolic Objectives and Tradeoffs in Single Cells During Embryogenesis

**DOI:** 10.1101/2024.02.09.579737

**Authors:** Da-Wei Lin, Ling Zhang, Jin Zhang, Sriram Chandrasekaran

**Affiliations:** Center for Bioinformatics and Computational Medicine, Ann Arbor, MI, 48109, USA; Department of Statistics, University of Michigan, Ann Arbor; Department of Biomedical Engineering, University of Michigan, Ann Arbor; Program in Chemical Biology, University of Michigan, Ann Arbor; Rogel Cancer Center, University of Michigan Medical School, Ann Arbor, MI, 48109, USA; Center for Stem Cell and Regenerative Medicine, Department of Basic Medical Sciences, and Bone Marrow Transplantation Center of the First Affiliated Hospital, Zhejiang University, Hangzhou, China; Liangzhu Laboratory, Zhejiang University, Hangzhou, 311121, China

## Abstract

While proliferating cells optimize their metabolism to produce biomass, the metabolic objectives of cells that perform non-proliferative tasks are unclear. The opposing requirements for optimizing each objective results in a trade-off that forces single cells to prioritize their metabolic needs and optimally allocate limited resources. To define metabolic objectives and tradeoffs in biological systems mathematically, we integrated bulk and single-cell omics data with a novel framework to infer cell objectives using metabolic modeling and machine learning. We validated this framework by identifying essential genes from CRISPR-Cas9 screens in embryonic stem cells, and by inferring the metabolic objectives of quiescent cells and during different cell-cycle phases. Applying this to embryonic cell states, we observed a decrease in metabolic entropy upon development. We further uncovered a trade-off between glutathione and biosynthetic precursors in 1-cell zygote, 2-cell embryo, and blastocyst cells, potentially representing a trade-off between pluripotency and proliferation.

## Introduction

Cell-type transitions, including embryogenesis, proceed via a cascade of changes by switching among different metabolic pathways (Shyh-Chang and Ng 2017; Cliff and Dalton 2017). This complex metabolic rewiring can make it challenging for cells to optimize competing biological objectives (Hausser et al. 2019). This is explained by the concept of Pareto optimality and the cost-benefit theory which describes the trade-off in multiple metabolic objectives (Wortel et al. 2018). Prior studies have found that enzyme cost, respiratory activity, and growth rate form a Pareto space in *E. coli* in which the three objectives compete (Schuetz et al. 2012). Using gene expression data, prior studies have uncovered evolutionary trade-offs in exponential and stationary phase gene activity in microbes (Shoval et al. 2012). Nevertheless, metabolic trade-offs in multicellular systems and cell-state transitions have not been evaluated yet (Kuzawa et al. 2014; Korem et al. 2015; Hausser and Alon 2020).

Although theoretical models such as flux balance analysis can be used to assess the intricacies of these transitions, cell growth is usually assumed to be the objective in these models (O’Brien et al. 2015) (Bordbar et al. 2014). Traditional assumptions of maximizing biomass that is commonly used in the modeling of metabolic networks do not apply to all living systems (Bordbar et al. 2014). For example, dormant cells or terminally differentiated mammalian cells do not maximize biomass. Further, during certain embryonic cell stages, no significant growth is observed. Early embryos likely need to alter their metabolic objectives to transition between growth stages. Particularly in the transition from zygote/1-cell (1C) to 2-cell (2C) stage, the dependence on maternal materials is reduced and zygotic genome activation (ZGA) is initiated. The metabolite pyruvate fuels the transition and activates epigenetic processes in the nucleus (Nagaraj et al. 2017). On the other hand, the transition from 2C to the blastocyst (BC) stage involves the differentiation of cell types and increases in cell numbers from 2 to 32. Prior studies based on the transcriptome, epigenome, and metabolome suggest up-regulation of respiration and higher consumption of glucose in this transition (J. Zhao et al. 2021).

How embryos optimize their metabolism and allocate resources for dramatic changes from one stage to another is unclear. Phenotypically, the two transitions double cell numbers, but the total volume is almost consistent (Chen, Qian, and Good 2020). In fact, as we mentioned previously, the top priority tasks are ZGA and differentiation in 1C2C and 2CBC, respectively. Optimization of biomass production alone is hence unlikely to explain the metabolic rewiring in embryogenesis. Indeed, development rather than growth can be an alternative goal for embryos.

In addition to embryos, most other mammalian cell types may not optimize their metabolism for biomass synthesis (Bordbar et al. 2014). For example, quiescent cells phenotypically stop increasing their biomass. Even in proliferating cells, the G1 and S phases in the cell cycle have been known to have two distinct metabolic functions, the accumulation of amino acid and the production of nucleotides, respectively (Kalucka et al. 2015). Although several methods have been developed to infer biomass composition in microbial strains, there is no consensus on suitable objective functions in mammalian cells ((Devoid et al. 2013); (National Research Council et al. 2015); (National Research Council et al. 2015; Gianchandani et al. 2008); (Q. Zhao et al. 2016); (Lachance et al. 2019)). Instead of directly inferring objective functions, two recently developed methods alternatively identified metabolic tasks in mammalian and cancer cells from omics datasets ((Richelle et al. 2021) and (S. Gao et al. 2022)). Yet these pioneering studies still require non-zero fluxes of biomass reactions, and do not account for the trade-offs in optimizing various metabolites.

To investigate the goals of metabolic rewiring in cell-type transitions, here we developed a computational framework based on optimization theory to infer cell-specific metabolic objectives called Single Cell Optimization Objective and Tradeoff Inference (SCOOTI). We demonstrate that this computation framework can identify metabolic objectives of proliferative and quiescent cells, and during different cell-cycle phases based on both bulk and single-cell transcriptomics datasets. We then applied our approach to decipher metabolic objectives and trade-offs in mouse early embryogenesis.

## Results

### SCOOTI to infer cell-specific metabolic objectives

To define metabolic goals of a cell mathematically, we develop a novel theoretical framework to infer cellular objectives from omics data. By integrating metabolic models with omics data under various conditions our approach identifies metabolic objectives in different cell types and cell-type transitions. The underlying hypothesis for our approach is that cells may optimize a small subset of biomass components at any given time. For example, there is a division of labor in metabolic tasks over the cell cycle phases, and single cells do not maximize all biomass components simultaneously (Birch, Udell, and Covert 2014). The output from our approach is a set of non-negative weights corresponding to the importance of each biomass component in a given condition.

Firstly, a set of optimal metabolic flux vectors is determined by individually optimizing the production of 52 different metabolites that are typically used in biomass objectives (Table 1). This list includes precursors and cofactors involved in energy production, redox status, precursors for metabolic and epigenetic processes, and reducing stress. Secondly, omics-constrained models are used to determine metabolic fluxes that are most consistent with up- and down-regulated metabolic enzymes or metabolites during cell type transitions (Figure 1). Finally, meta-learner regression is used to determine the combination of optimal flux vectors that best represent the flux solution for a given cell state (Methods). The meta-learner regression model was trained with optimal flux vectors as features and constrained models as outcomes. This results in a list of regression coefficients, corresponding to each of the 52 optimal flux vectors, that best describe the metabolic state from omics data. Representing a system through these metabolic objectives could be considered as a feature reduction process wherein a combination of 52 optimal flux vectors summarizes the information from thousands of context-specific reaction fluxes.

**Figure 1.**
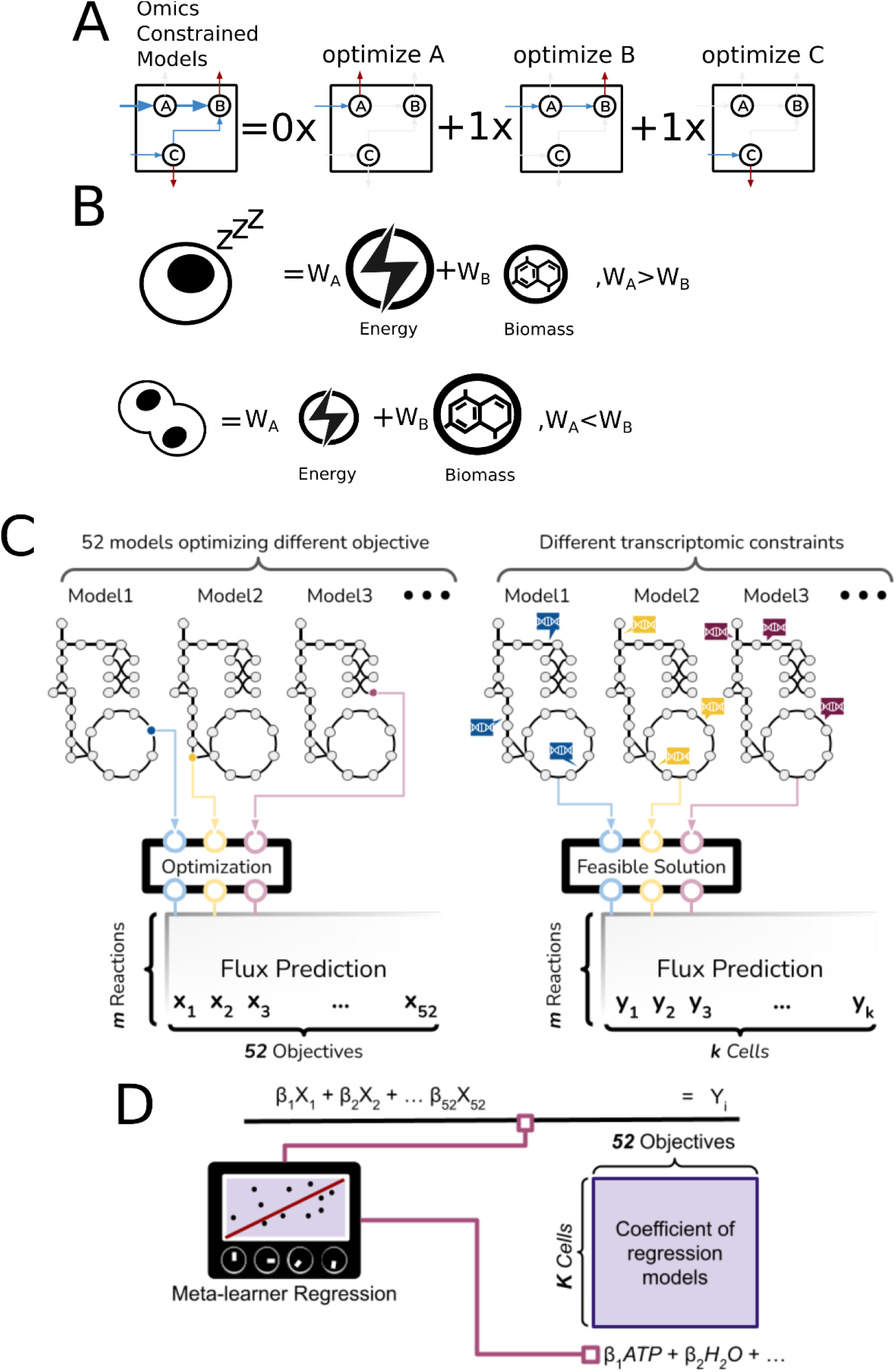
Using SCOOTI to infer cell-specific metabolic objectives. (A) A simplified example of a regression model learned from metabolic models optimizing single metabolites A, B, and C (right-hand side) to predict the omics-constrained models (left-hand side). The models optimizing B and C together can approximate the omics-constrained model, suggesting higher demand fluxes for B and C. (B) Cellular phenotypes are supported by different metabolic objectives. For instance, cell quiescence (Top) is assumed to depend more on energy metabolism than biomass. In contrast, proliferating cells may have lower demand on energy metabolism compared to biomass (Bottom). (C) The presented computational framework, SCOOTI, requires two types of models. (1) Genome-scale Metabolic Models (GEMs) without omics constraints were used to predict the optimal metabolic flux allocations to maximize the production of 52 different metabolites individually using flux balance analysis (Left). (2) Omics-constrained models without objective functions were used to determine optimized flux allocations given up- and down-regulating metabolic genes/proteins/metabolites according to cell- or cell-type-specific omics data (Right). (D) Meta-learner regression models, which comprise 4 different regression models, regressed 52 unconstrained models on the constrained models. The coefficients suggested linear combinations of 52 metabolites that formulated cell- or cell-type-specific metabolic objectives.

**Table 1.** Metabolites and their corresponding ID in the GEM.

| Amino acids |  |  |  | Nucleotides |  | Lipid |  | Phospholipid |  | Carbohydrate and other molecules |  |
| --- | --- | --- | --- | --- | --- | --- | --- | --- | --- | --- | --- |
| Metabolite | BIGG ID | Metabolite | BIGG ID | Metabolite | BIGG ID | Metabolite | BIGG ID | Metabolite | BIGG ID | Metabolite | BIGG ID |
| Alanine | ala-L | Serine | ser-L | ATP | atp | Diacylglycerol | dag_hs | Phosphatidic acid | pa_hs | Glycogen | glygn1 |
| Arginine | arg-L | Threonine | thr-L | AMP | amp | Triacylglycerol | tag_hs | Phosphatidyl inositol | pail_hs | H2O | h2o |
| Asparagine | asn-L | Tryptophan | trp-L | CMP | cmp | Monoacylglycerol 2 | mag_hs | Phosphatidyl choline | pchol_hs | Biomass | gh |
| Aspartate | asp-L | Tyrosine | tyr-L | GMP | gmp | Cholesterol ester | xolest_hs | Phosphatidyl ethanolamine | pe_hs | Acetyl-CoA | accoa |
| Glutamine | gln-L | Valine | val-L | UMP | ump | Cholesterol | chsterol | Phosphatidyl serine | ps_hs | S-Adenosylmethionine | amet |
| Glutamate | glu-L | Methionine | met-L | dAMP | damp |  |  | Sphingomyelin | sphmyln_hs | Reduced glutathione | gthrd |
| Glycine | gly | Cysteine | cys-L | dCMP | dcmp |  |  | Cardiolipin | clpn_hs | Oxidative Glutathione | gthox |
| Histidine | his-L | Phenylalanine | phe-L | dGMP | dgmp |  |  | Lysophosphatidylcholine | lpchol_hs |  |  |
| Isoleucine | ile-L |  |  | dTMP | dtmp |  |  |  |  |  |  |
| Leucine | leu-L |  |  | NADH | nadh |  |  |  |  |  |  |
| Lysine | lys-L |  |  | NADPH | nadph |  |  |  |  |  |  |
| Proline | pro-L |  |  | NMN | nmn |  |  |  |  |  |  |

### Benchmarking the framework by inferring metabolic objectives of proliferative and quiescent cells

As a proof-of-concept, we applied SCOOTI to investigate proliferative and quiescent cells as they clearly have distinct metabolic objectives. We collected three bulk transcriptomic datasets that measured multiple types of growth stimulation and multiple types of quiescence induction, including contact inhibition, serum starvation, and cytokine starvation to validate our method (Min and Spencer 2019; Mitra et al. 2018; Sharma et al. 2021). To quantify the differences from the commonly assumed metabolic goal of maximizing biomass, we measured the Euclidean distance between the objective coefficients inferred from omics data and the coefficients using the biomass as objective. Coefficients from proliferative cells were much closer to the biomass objective than quiescent cells (Figure 2B). We next determined which metabolites among the 52 in our model best differentiate the two cell states, by statistically comparing the regression coefficients in quiescent and proliferative cells. The coefficients for glycine, nicotinamide mononucleotide (NMN), and leucine were significantly higher in proliferative cells while no metabolite was significantly greater in quiescent cells (Figure 2D). Visualizing the candidate metabolites selected in each of the transcriptomics-based objective revealed that cholesterol, NMN, glycine, and tyrosine were required for more proliferative states (Figure 2F, S3A-C). Although the coefficients were not significantly different from proliferative cells, lysophosphatidylcholine, alanine, and reduced glutathione were chosen by at least 50% of quiescent cell states (Figure 2F).

**Figure 2.**
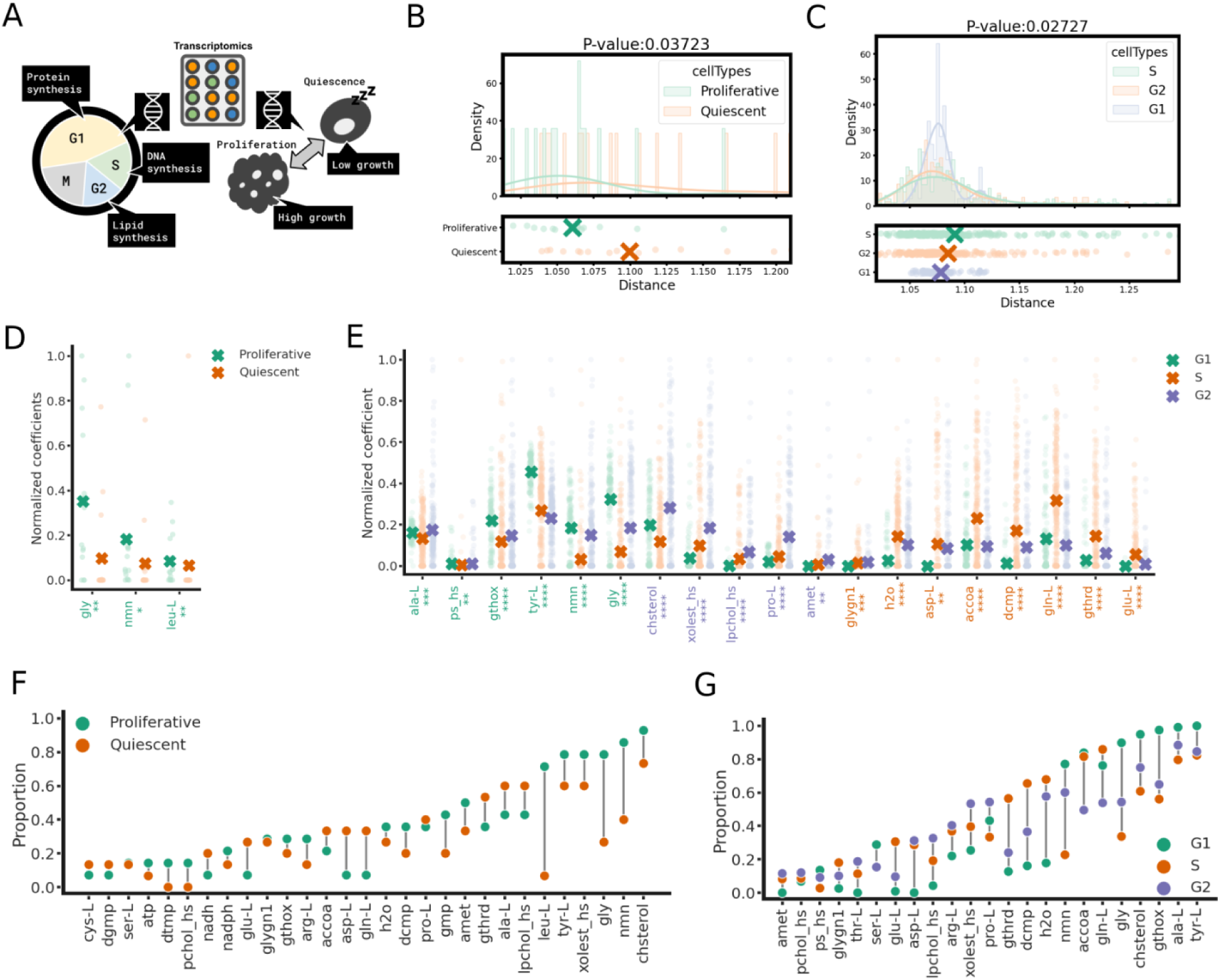
Inference of metabolic objectives in different cell types with known metabolic behaviors. (A) Graphical summary of known metabolic functions in cell-cycle phases and cell quiescence, such as high protein synthesis found in G1, high DNA synthesis found in S, and high lipid synthesis found in G2. Particularly, G1 and G2 have faster growth rates (Icard et al. 2019; Cheeseman 2020). (B, C) Euclidean distances of coefficients between the biomass objective and inferred metabolic objectives. (D) Statistical tests identified metabolites including glycine, NMN, and leucine with coefficients of metabolic objectives significantly higher in proliferative states. (E) Metabolites which had significantly greater coefficients in G1, G2, and S phases were marked green, orange, and purple on the x-axis, respectively. (F) Metabolites selected by a significant proportion of cells (> 10%) were shown in this plot. Cholesterol was the most selected metabolite in both states, but Leucine has the greatest difference which was chosen by less than 10% of quiescent cells but around 70% of proliferative cells. (G) Tyrosine and alanine were chosen by over 80% of cells in all the phases. In contrast, glycine, glutamine, and proline show highest proportions in the G1, S, and G2 phases, respectively. (D, E) The metabolic coefficients with p-values smaller than 0.05, 0.01, 0.001, and 0.0001 were labeled with *, **, ***, and ****, respectively.

Interestingly, the metabolic objectives in the proliferative cells mostly consist of growth-boosting metabolites: glycine supports muscle cell proliferation (Jain et al. 2012; C. Lin et al. 2020), stem cell renewal (Shyh-Chang and Ng 2017), and cancer cell growth (Jain et al. 2012), while leucine is associated with protein metabolism, proliferation, and inhibition of apoptosis (Gonçalves and Gomes-Marcondes 2010; Mao et al. 2013; Coëffier et al. 2011; Xiao et al. 2016). Additionally, NMN is a well-known anti-aging metabolite (Mills et al. 2016), and is reported to restore cells from senescence, promote proliferation, and increase cell viability (Yamaura et al. 2022; Pu et al. 2022; Fang et al. 2021). These results suggest that our model can infer metabolic objectives and distinguish cells with different proliferation abilities.

### Our framework recalls known metabolic division of labor during the cell cycle

We next applied SCOOTI to infer metabolic objectives using single-cell RNA sequencing (scRNA-seq) data to determine metabolic objectives during different cell-cycle phases in HeLa cells (Schwabe et al. 2020). Identifying cell-specific metabolic objectives could guide us to identify the heterogeneity of metabolic goals and intermediate cell types. In the comparisons between G1, S, and G2 stages, the metabolic objectives of G1 was closest in terms of Euclidean distance from the biomass objective (Figure 2C). Growth-associated metabolites, such as tyrosine, alanine, glycine, and NMN, were significantly higher (Figure 2E) and were chosen by a higher proportion of cells in the G1 phase (Figure 2G), consistent with protein synthesis and increased cell volume in this phase. By contrast, significantly higher dCMP, aspartate, and glutamine in the S phase is consistent with DNA synthesis in this phase (Figure 2E). Greater weights for cholesterol and lysophosphatidylcholine suggest an increase in lipid synthesis in G2 (Icard et al. 2019; W. Lin and Arthur 2007). In addition, higher reduced-glutathione in the S phase suggested a potential need for protection from oxidative stress in this phase (Figure 2E). Glutathione has been observed to accumulate in the nucleus during the S and G2 phases before mitosis (Markovic et al. 2007). Reduced glutathione concentration reached its peak at the S phase although the oxidized form peaks at the G1 phase (Markovic et al. 2007; Ahn et al. 2017). This is consistent with significantly greater oxidized-glutathione coefficients in the metabolic objectives of G1. Consistent with the differences in metabolic objectives between G1, S, and G2, cells in these phases were clearly separated with Uniform Manifold Approximation and Projection (UMAP) (Figure S4A). Interestingly, the UMAP of objectives and reconstructed fluxes showed better separation of cell states than the transcriptomics- and flux-based UMAP (figure S8A-D, top).

### Validation of the objective-inference framework using CRISPR-Cas9 screens in stem cells

We next validated the model by its ability to identify essential genes in mouse embryonic stem cells (mESC) based on CRISPR-Cas9 screens (Shohat and Shifman 2019; Tzelepis et al. 2016). We hypothesized that ESCs have non-canonical metabolic objectives, such as maintaining pluripotency instead of purely maximizing biomass. We first inferred metabolic objectives using transcriptomics data for wild-type mESCs (Y. Zhang et al. 2013) We then performed *in silico* gene knockouts and considered a gene as essential if it led to a significant change between wild-type and knockout metabolic objectives (Figure S1A and B). This approach outperformed traditional biomass objectives in predicting gene essentiality across a wide range of parameters (Figure S1C). Notably, genes correctly identified as essential by our framework were enriched in Gene Ontology (GO) terms associated with Oxidative Phosphorylation (see Figure S10A and C). This is consistent with the connections between the Oxidative Phosphorylation and early pluripotency (Zhou et al. 2012). In contrast, the traditional biomass method primarily identified genes related to biosynthetic processes, resulting in several false positives. Thus, the objective-inference approach has the potential to uncover gene essentiality in cellular phenotypes beyond just cell growth. Overall, these results suggest that the inferred metabolic objectives can quantitatively summarize the metabolic goals of a cell.

### Transitions during early embryogenesis were driven by distinct metabolic objectives

To identify metabolic objectives using our framework in the two cell-type transitions, zygote to 2-cell (1C2C) and 2-cell to blastocyst (2CBC), during embryogenesis, we integrated gene expression levels, protein abundance, and metabolite concentrations with genome-scale metabolic models (Figure 3A). Specifically, based on transcriptomic or proteomic datasets, metabolic genes or enzymes that were significantly up- and down-regulated were determined by comparing the expression levels in 1C to those in 2C. Likewise, the concept was also applied to the transitions from 2C to BC. By doing so, the genes and enzymes were used to constrain the reaction fluxes. In addition, time-course metabolomic data were converted into additional constraints, and constrained GEMs were used to generate fluxes (Chandrasekaran et al. 2017; Shen et al. 2019). Meta-regression was then employed to predict the fluxes from constrained models as response with the 52 different optimal flux vectors as predictors, resulting in coefficients for the 52 metabolic objectives.

**Figure 3.**
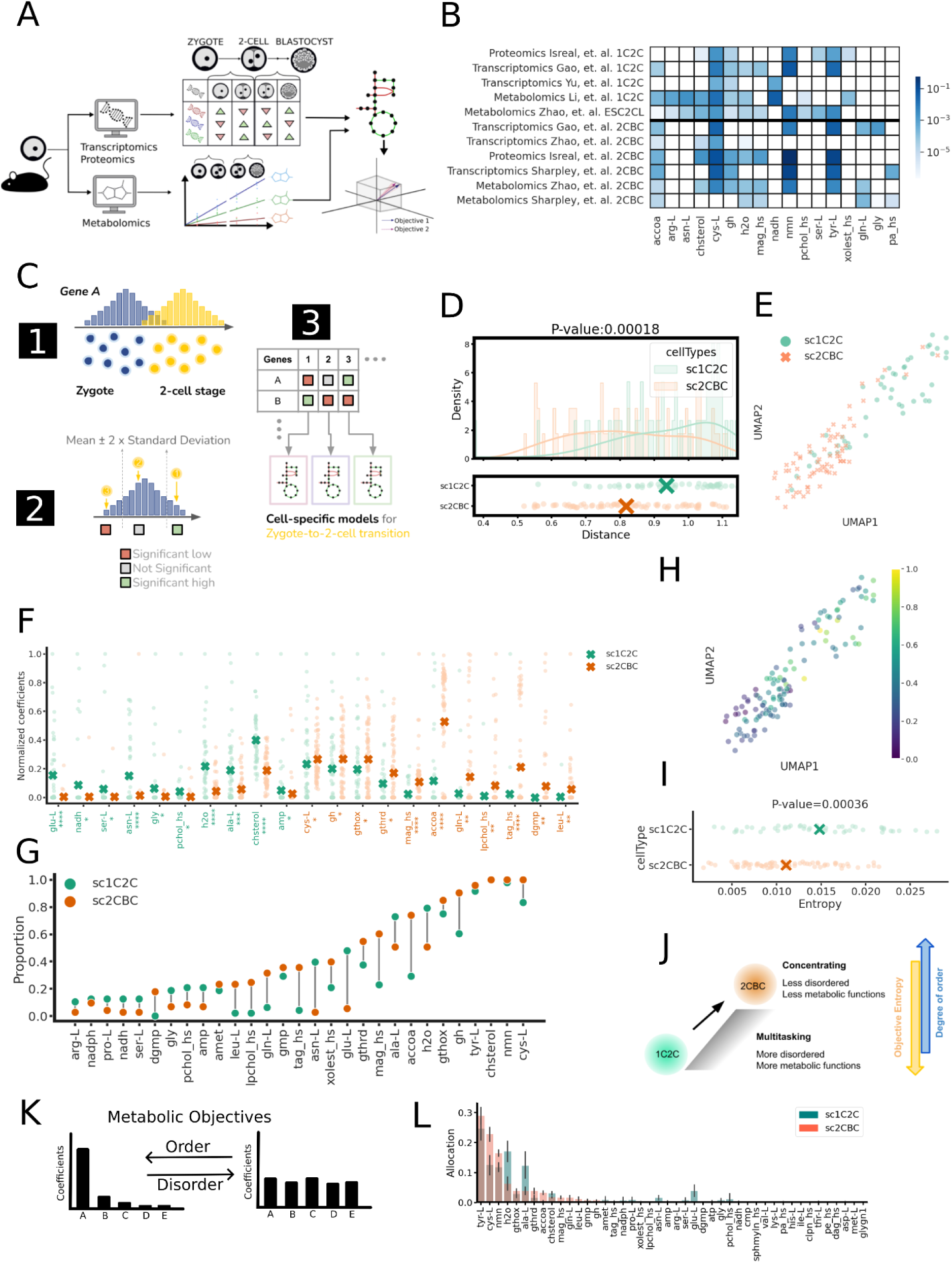
Metabolic objectives inferred in 1C2C and 2CBC transitions. (A) The pipeline for modeling flux allocations with mouse embryo multi-omics datasets. Either significant metabolic genes identified from transcriptomics or enzymes identified from proteomics were applied as constraints to infer fluxes (top). Time-course metabolomics data were converted into metabolite rate of change and used to infer fluxes (bottom). (B) Coefficients of metabolic objectives inferred with bulk multi-omics datasets of transitions from 1C to 2C and from 2C to BC. (C) Statistical method used to compute cell-specific constraints on single-cell embryogenesis datasets. (D) Euclidean distance from the Recon1’s biomass objective function to cell-specific metabolic objectives indicates significantly closer coefficients in the 2CBC transitions. (E) UMAP reduced the dimension of metabolic objectives of the two transitions and was visualized as a scatterplot. (F) Statistical tests identified the biomass precursor (gh), acetyl-CoA, glutathione, and so on marked red were significantly higher in the 2CBC transitions. Other metabolites including cholesterol, alanine, and glycine were significantly higher in the 1C2C transitions. Significant metabolites with p-values smaller than 0.05, 0.01, 0.001, and 0.0001 were labeled with *, **, ***, and ****, respectively. (G) Cysteine, NMN, cholesterol, and tyrosine were selected by more than 80% of both transitions as metabolic features. However, the biomass precursor was relatively greater in the 2CBC transitions. (H) Entropy of the objectives overlaid on the UMAP visualization showed an increased trend from the bottom to the top. (I) Entropy of the objectives was significantly greater in the 1C2C transitions. (J) The illustration explained the relationship between objective entropy and the metabolic goals. The objective entropy can be linked with the allocation of metabolic functions. Higher entropy implies a multitasking metabolic system. Entropy represents the degree of disorder in a system. A simplified example of metabolic system (K, left) relying on fewer metabolic objectives (more ordered, less entropy) thus showing greater coefficients for a few metabolites. In contrast, disordered metabolic objectives in another simplified example (K, right) has a greater spread in coefficients of metabolites, which implies a network optimized for multitasking. (L) The 1C2C transition with significantly greater entropy than the 2CBC transition shown in (I) was confirmed by the allocation of coefficients. The 1C2C and 2CBC transitions showing multitasking and concentrating allocations, respectively, can map to the two simplified examples shown in (K).

NMN, tyrosine, and cysteine consistently had the largest coefficients of metabolic objective in both 1C2C and 2CBC transitions based on metabolomics, transcriptomics, and proteomics (Figure 3B). Oxidized and reduced glutathione and S-adenosyl methionine (SAM) were also identified as important features based on transcriptomics and proteomics (Figure S3D). Analysis of metabolomics and proteomics data suggest that cholesterol and acetyl-CoA were important for both 1C2C and 2CBC transitions. In contrast to these common metabolic objectives, glutamine was present in the 2CBC transition while glutamate, was significant only in the 1C2C transition. It is noteworthy that the biomass objective (gh), which consists of all metabolites used in the human metabolic model’s biomass objective has a greater magnitude of coefficient in the 2CBC transition. The optimal flux vector optimizing biomass was included in our model as an additional control feature.

Due to the stochastic nature of regulatory networks in embryos, metabolic objectives may vary from cell to cell. To address this and quantify the heterogeneity of metabolic objectives during embryogenesis, we integrated single-cell transcriptomics datasets into SCOOTI (Figure 3C). Similar to the analysis of cell-cycle phases (Figure 2E), the regression coefficients were statistically tested between 1C2C and 2CBC transitions.

The biomass objective showed significantly high coefficients in the 2CBC transition directly indicating higher growth rate (Figure 3F). This was supported by significantly high glutamine and leucine coefficients (Figure 3F, 3G). Glutamine is associated with cell proliferation (Xi et al. 2011), redox homeostasis (Amores-Sánchez and Medina 1999), and initiates early differentiation by α-Ketoglutarate (TeSlaa et al. 2016). The association between leucine and cell proliferation is also observed in T-cells (Ananieva, Powell, and Hutson 2016), fibroblasts, (Gonçalves and Gomes-Marcondes 2010) and embryonic stem cells (Correia et al. 2022). These three metabolites, because they are associated with growth, point towards greater proliferation in the 2CBC transition, which aligns with the bulk-omics data models (Figure 3B). The finding was also confirmed by comparing the inferred embryogenesis objectives to the traditional biomass objective function. The Euclidean distance of coefficients in the cells undergoing the 2CBC transition was closer to the coefficients of the biomass objective in the human metabolic reconstruction, which implies more growth and biomass production in this transition (Figure 3D).

As for the 1C2C transition, glutamate, glycine, and asparagine were identified as significant and selected by about 50% of the cells in this transition (Figure 3F, G). In fact, the turnover rates of these three amino acids changed significantly in Day 2 cryopreserved cleavage-stage compared to Day 1 one-cell cryopreserved human embryos (Picton et al. 2010). These three metabolites play an important role in this phase. For instance, low glutamine-to-glutamate ratio can reduce DNA fragmentation (Drábková et al. 2016) while glycine promotes nucleotide synthesis (Tedeschi et al. 2013); these align with the observation of DNA replication and the initiation of transcription (ZGA) during the 1C2C transition (Jukam, Shariati, and Skotheim 2017; Lee, Bonneau, and Giraldez 2014) (Figure 3F, G). Cholesterol and cysteine also showed significant differences and were higher in the 1C2C and the 2CBC transitions, respectively, although these two metabolites had relatively high regression coefficients for both states (Figure 3G). Monoacylglycerol, triglyceride, LPC, and acetyl-CoA, however, showed significantly higher coefficients in the 2CBC transition and emphasized the high priority of lipid synthesis and energy storage (figure 3F). High proportion of glutathione in both transitions, based on scRNA-seq, suggested the essentiality of redox homeostasis up to the blastocyst stage (Winkler et al. 2011).

The single-cell metabolic objectives of 1C2C and 2CBC transitions were then projected onto a low-dimensional space by UMAP (Figure 3E). Interestingly, the UMAP of objective coefficients showed a linear trend from 1C2C to 2CBC transitions. We were inspired by the linear transitions on UMAP and hypothesized that the metabolic objectives may correspond to distinct cellular phenotypes (Figures 4E). To verify if the metabolic objectives are associated with embryonic development, enrichment analysis was performed for four gene sets related to pluripotency, development, growth, and proliferation. The results showed that the genes of ESC pluripotency and blastocyst development were significantly enriched in one of the two clusters comprising 1C2C cells (Figures 4F). Trophectoderm cell proliferation and blastocyst growth were enriched in the cluster dominated by the 2CBC transitions (Figure 4G). This suggests that changes in metabolic goals were associated with embryogenesis. Therefore, pluripotency/development and proliferation/growth could be mapped to the two metabolic traits in a two-dimensional representation of metabolic objectives (Figure 4F and G, bottom).

**Figure 4.**
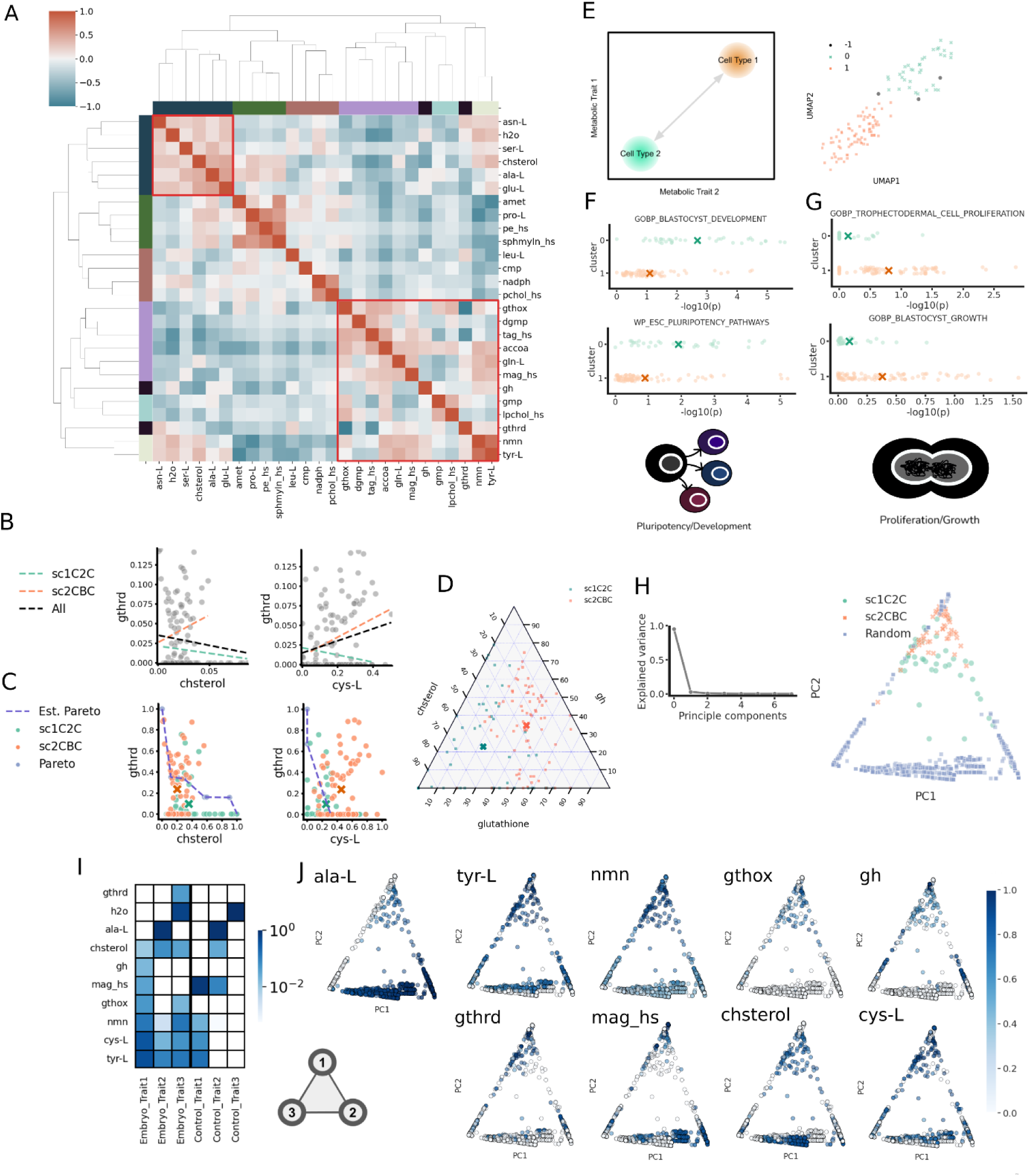
Trade-offs in metabolic objectives during embryogenesis. (A) Relationships among metabolites selected by metabolic objectives of embryogenesis were quantified with Pearson correlations. Correlation coefficients were normalized with the lowest values. Negative correlations implied potential trade-offs. (B) Scatter plots depict the relationships of objective coefficients among reduced glutathione, cholesterol, and cysteine. The dash lines represented the best-fit of coefficient relationships in different transitions. (C) Normalized objective coefficients (green and orange) overlaying Pareto fronts of ideal objective fluxes (blue) identified if the trade-offs between any pair of objectives were closer to the optimality. The means of coefficients were shown as ‘X’. (D) The proportion of coefficients that represent metabolite allocation among cholesterol, the biomass precursor, and the summation of oxidized and reduced glutathione were shown as percentages. (E) UMAP Clustering results with HDBSCAN showed two groups which were labeled 0 and 1 for the following analysis in panels F and G. The interpretation of this grouping corresponding to cell phenotypes is illustrated in the left panel. F and G. Functional enrichment analysis of gene sets in the two groups using GSEA MsigDB demonstrated that (F, top) blastocyst development and (F, middle) ESC pluripotency pathways were significantly enriched in group 0 while (G, top) trophectodermal cell proliferation and (G, middle) blastocyst growth were enriched in group 1. (F and G, bottom) H. A PCA plot automatically extracted 3 traits, according to the Elbow method based on the normalized ideal objectives. (I) The PCA trait plots were respectively overlaid with the allocation of the top 50% of candidate metabolites. J. The metabolic objectives (Embryo_Trait) corresponding to the three vertices are shown in a heatmap and compared against control traits obtained without any omics constraints suggesting that those are not embryo-specific and driven by inherent structure of the metabolic network.

### Metabolic trade-offs during early embryogenesis

Embryonic development, based on single-cell transcriptomics, is anticipated to reduce entropy and lead to an ordered transcriptomic structure (Waites and Davies 2019; Teschendorff and Enver 2017). To investigate the degree of disorder in metabolic objectives, we quantified how embryos deployed their metabolic tasks by measuring the entropy of the inferred objectives. The measurements showed a decreasing trend along the UMAP plot of objective coefficients (Figure 3H). Statistically, the objective entropy was significantly lower in the 2CBC transitions (Figure 3I). In other words, the cells undergoing 2CBC transition concentrated on fewer metabolic objectives (Figure 3J). The finding can be confirmed by the allocation of metabolite coefficients. Hypothetically, an ordered system may concentrate the coefficients in a few metabolites, but a system with high entropy in contrast, tends to weigh multiple metabolites similar levels of coefficients (Figure 3K). As a result, the allocation of coefficients indicated that more metabolites were weighted with comparable levels of coefficients in the 1C2C transition than in the 2CBC transitions (Figure 3L).

To investigate if trade-offs were shown in metabolic objectives during embryogenesis, we quantified the correlation between the regression coefficients of metabolites. We identified consistent pairs of metabolites that showed strong negative correlations which suggested a tradeoff (Garland, Downs, and Ives 2022). A subset of metabolites including both oxidized and reduced glutathione were more significant in the 2CBC transition (right bottom cluster of the heatmap; Figure 4A). These metabolites were negatively correlated with glutamate, alanine, cholesterol, and asparagine which were significant in the 1C2C transition (Figure 4A).

Since reduced glutathione has been reported as a crucial metabolite related to embryogenesis, pluripotency, and differentiation (Hansen and Harris 2015; Ufer and Wang 2011), we investigated how glutathione influences metabolic tasks during the two transitions with Pareto analysis. According to the theory of optimality, tradeoffs in objectives can lead to optimal conditions lying on Pareto fronts, which represents the best solutions for optimizing multiple objectives (Nagrath et al. 2007; Schuetz et al. 2012). Any points inside the region formed by Pareto fronts are feasible but not optimal solutions; outside the region, however, the points are not biologically feasible. Using this idea, we modeled ideal Pareto fronts with unconstrained models and by randomizing objective coefficients. By doing so, we can inspect if there were any pairs of candidate metabolites following the inherent limitations in the metabolic network, so-called mechanistic trade-offs (Weiße et al. 2015; Hashemi, Laitinen, and Nikoloski 2023).

The average coefficients of both transitions were found to be close to Pareto fronts formed by cholesterol and reduced glutathione (Figure 4C, left). Interestingly, the Pareto front of cysteine and reduced glutathione passed through the average coefficients of the 1C2C but not the 2CBC transition (Figure 4C, right). Given that cysteine and cholesterol were common features (selected by more than 80% of single-cells) in the two transitions, the result above suggests that redox homeostasis was optimized and required during embryogenesis. We further examined the relationships between glutathione and other metabolites with scatter plots. Surprisingly, reduced glutathione only showed negative relationships, implying a trade-off, only in the 1C2C transition while the two pairs became positive relationships in the 2CBC transition (Figure 4B). Glutathione, cysteine and cholesterol have been reported to influence early pregnancy, fertilization rate, and survival of early embryo (Anchordoquy et al. 2019; Baardman et al. 2013; Shi et al. 2000). Given that the 1C2C transition heavily relies on maternal sources and more active protein synthesis and metabolic activities after ZGA, these results imply that embryo mitigate the trade-offs in the 2CBC transitions (Jukam, Shariati, and Skotheim 2017).

The negative relationship between the biomass objective and glutathione in the 1C2C transition (Figure S6D, S6E) suggests that redox functions are competing with embryo growth in this stage (Hansen and Harris 2015). However, the 2CBC transition did not show the negative relationship. The first evidence supporting this observation is a greater drop of reduced glutathione in the transition from oocyte to 2C than the transition from 2C to BC (Gardiner and Reed 1994). Reduced glutathione shows relatively less change in the 2CBC transition and leads to a more oxidative status in BC (J. Zhao et al. 2021) hinting at a lower need of protection from redox stress. The second evidence is that BC but not 2C can replenish reduced glutathione (Gardiner and Reed 1995).

Additionally, a notable shift from the 1C2C to the 2CBC transition was from low to high allocation of the biomass objective. Interestingly, cholesterol also competes with glutathione and biomass in both transitions (Figure 4D). The essentiality of cholesterol has been reported in mouse embryo (Wolf 1999). Removing cholesterol led to abnormality in rat embryo (Roux et al. 2000). Since cholesterol is a structural component for cell membrane, the trade-off can be explained when embryo growth requires both the production of biomass precursor and extensive synthesis of cell membrane.

We next explored high dimensional trade-offs in metabolic traits during embryogenesis by analyzing groups of metabolites. We selected the metabolites that were chosen by at least 50% of each transition to construct ideal metabolic traits (Figure 3G). Employing Principal Component Analysis (PCA) to project these allocations into a three-dimensional space, the ideal allocations shaped the boundaries of a polygon. This guided us in extracting three components with a high explained variance ratio. PCA of the proportion of metabolic objectives identified three competing metabolic traits (Figure 4H). Each vertex of the triangle represents an ideal metabolic trait. Oxidized glutathione, the biomass objective, and cholesterol were closest to the top vertex, which predominantly contained cells in 2CBC transition (Figure 4I). NMN, cysteine, tyrosine, and cholesterol were high in the bottom two vertices. NMN is known for its functions in the homeostasis of NAD^+^ and potential anti-aging effects in tissues and embryos. We found that pyrimidine and IMP biosynthesis were the top metabolic pathways associated with the top vertex. This supported the longer distances of the cells in 1C2C transition, characterized by lower growth rates and transcriptional activities due to ZGA, when compared to the 2CBC transition. Additionally, the demand for fatty acids (Xiang et al. 2020) and pyruvate during ZGA (Sharpley et al. 2021; T. Zhang et al. 2022) resulted in cells from the 1C2C transition cluster towards the second and the third vertices in the bottom of the triangle, where fatty acid elongation and pyruvate metabolism were the top metabolic pathways.

To assess if these trade-offs are universal or context-specific, we predicted metabolic trade-offs during the cell cycle (Figure 2E). Correlation analysis among the objectives identified 3 sets of metabolites that have relatively strong trade-offs to other sets (Figure S9A). Notably, a distinct trade-off emerged between glutathione, dCMP and cholesterol, representing these 3 distinct clusters of metabolites (Figure S9B). Pareto analysis highlighted the interplay among redox, nucleotide, and lipid metabolism, shaping the metabolic characteristics of cell-cycle phases, with corresponding metabolites exhibiting significantly greater coefficients in G1, S and G2 phases respectively (Figure S9C). Thus, the tradeoffs between glutathione and biosynthetic processes may shape the metabolic behavior in other biological processes as well.

## Conclusion

To define the metabolic goals and tradeoffs during cellular processes mathematically, here we develop a novel theoretical framework to infer cellular objectives from both omics data and knowledge of metabolic network architecture. Traditional assumptions of maximizing biomass that are commonly used in modeling of metabolic networks do not apply to all living systems. The biomass objective, consisting of amino acids, nucleotides, phospholipids, and other essential metabolites, represents the rate of growth. However, cells may not produce all these biomass components simultaneously and the utility of biomass objectives has been questioned for mammalian systems.

In this study, we propose a novel approach that involves data-driven inference of metabolic objectives, consolidating information from numerous transcriptomics datasets and more than 3,000 metabolic reactions into 52 condition- or cell-specific features. We use meta-regression to determine the combination of optimal flux vectors that best represent the metabolic objective for a given cell state. Our results suggest that cellular phenotypes can be quantified through linear combinations of metabolic objectives. Further, this study demonstrates that important metabolic functions can be recapitulated by metabolic objectives and used to separate cell types in low-dimensional spaces with both bulk and single-cell datasets. Of note, metabolic objectives are not necessarily the same as metabolic states. Different metabolic states, i.e. activity of various reactions in the network, can lead to the same metabolic objectives. Also, a high objective for a metabolite in a condition may not result in higher absolute levels for that metabolite and may instead represent a higher demand or turnover of the metabolite.

Our approach has some limitations related to the use of genome-scale metabolic models and machine learning. The quality of the metabolic models, the dataset used, and the method of data integration may strongly influence predictions (Gu et al. 2019). Nevertheless, we found that the results are robust over a wide range of parameters and thresholds used for integration of omics data (Figure S1) and similar results were obtained using different data sources and data types (Figure S3A-C), suggesting that our approach is able to uncover strongest biological signals from the data. The use of 52 candidate objectives, while comprehensive, may not completely capture objectives involving higher order effects like production of specific proteins. There may also be metabolites that are highly correlated and may be chosen interchangeably by machine-learning methods. The use of parsimonious methods like Lasso in our approach helps mitigate the issue of correlated variables.

Using this novel approach, we identify metabolic objectives in different cell types and cell-type transitions, including cell quiescence and cell-cycle. Applying this approach to single cell transcriptomics data from HeLa cells undergoing cell cycle correctly revealed metabolic objectives in each phase. In addition, metabolic objective trade-off analysis suggests that this division of labor during the cell cycle may be due to inherent limitations in the metabolic network, as lipid, redox and nucleotide synthesis compete with each other and cannot be optimized simultaneously. This mirrors previous work in microbes that have shown that ATP yield and biomass production cannot be optimized at the same time (Schuetz et al. 2012). Applying this approach to various types of quiescent cells revealed that their metabolic objectives are further away from biomass synthesis than proliferative cells, as expected. Interestingly, redox and lipid metabolism are active in quiescent cells, consistent with findings from prior studies that some core metabolic activities are maintained during quiescence.

Applying our tool to embryonic development unveiled the importance of glutathione and its critical role in the protection of oxidative stress during development. Optimizing the biomass precursor or growth-related metabolites such as glycine and NMN would be sufficient for cell proliferation but may not support development. This tradeoff is especially relevant during the first steps of development, where cells decide between proliferation, maintaining pluripotency, and differentiation. Our analysis suggests that glutathione, the biomass objective and associated metabolites like cholesterol potentially represent a Pareto optimality trade-off. These two metabolic objectives represent the contrasting choice between a state of pluripotency and proliferation, respectively. While it is known that the redox balance affects embryo phenotypes, here we uncovered the complex relationships among glutathione and biomass precursors and identified distinct stage-specific trade-offs. These trade-offs may play a crucial role in embryonic development and the maintenance of cell states. Especially during early embryogenesis, limitations in maternal resources and low metabolic and transcriptional activities may give rise to such trade-offs. Further research is necessary to determine whether perturbing such metabolic trade-offs can influence embryo phenotypes. Interestingly, both Pareto analysis and entropy analysis suggested that multiple metabolic goals are required in the 1C-2C early pluripotent state compared to later development with more deterministic cell fates.

Our study reveals the design principles and constraints in metabolic network remodeling during embryogenesis and other cellular processes. Notably, our approach bridges metabolic modeling and evolutionary biology and sheds light on metabolic trade-offs in these systems. Aside from being a hypothesis-generating tool, our approach can be beneficial for drug development and bioengineering applications. Trade-offs between metabolites could be used to selectively eliminate or enrich a certain cell type and enable evolutionary medicine or regenerative medicine applications.

## Methods

### Differential expression analysis

To identify differentially expressed genes in omics data for integration into the metabolic model, two different approaches were used, namely fold change and z-score, for bulk and single-cell data respectively. Significantly expressed genes in bulk cell quiescence datasets were calculated by comparing the treated and untreated conditions with T-test. We used 0.05 as a cutoff for p-values. Those significant genes were further classified into up- or down-regulated genes with fold-change higher than 1 or lower than 1. The same criteria and statistical methods were applied to bulk transcriptomics and proteomics datasets of embryogenesis. For the z-score approach for single-cell transcriptomics comparison of embryonic cell states, each reaction was z-score normalized across conditions. Here, transcripts with z-scores above or below a threshold of +2/-2 were considered upregulated or downregulated, respectively. For the cell cycle data, genes with expression level higher than the 40^th^ or lower than the 60^th^ expression level in each cell are considered up- or down-regulated genes, respectively. The thresholds were determined based on clear separation of groups on a UMAP plot.

### Integration of omics datasets into constraint-based metabolic models

The up- and down-regulated genes were integrated with a human GEM adapted from Recon 1 model with epigenetic reactions (Shen et al. 2019; Duarte et al. 2007). To incorporate the differentially expressed gene sets into the metabolic model, a linear version of the iMAT algorithm was used (Shen, Cheek, and Chandrasekaran 2019; Shen et al. 2019). In this approach, reactions associated with upregulated genes are assigned higher flux values. Reactions associated with downregulated genes are assigned lower values rather than being removed in contrast to the original iMAT approach. The iMAT approach can be used without any objective thus allowing for unbiased exploration of optimal objectives that best fit the omics data (Zur, Ruppin, and Shlomi 2010). To determine optimal input parameters for modeling, kappa (weight for downregulated genes), and rho (weight for upregulated genes), different ranges of values were tested to determine the best parameter set that separates cell types on UMAP. There were three steps in our criteria. Firstly, UMAP was used to reduce the dimension of flux data (McInnes, Healy, and Melville 2018). Secondly, we evaluated clustering performance by Silhouette scores and picked the combination of the parameters with the highest scores. If all the scores were negative, the low-dimensional representations were further clustered by Hierarchical Density-Based Spatial Clustering of Applications with Noise (HDBSCAN) and evaluated by mutual information and rand index (Campello, Moulavi, and Sander 2013). Constrained models for cell quiescence were generated with hyperparameter kappa 0.1 and rho 10, HeLa cell cycle for kappa 0.01 and rho 0.01, bulk multi-omics embryogenesis for kappa 0.01, rho 10, and and single-cell embryogenesis for kappa 0.1 and rho 0.01. Incorporating time-course metabolomics data of embryogenesis requires another pipeline (Chandrasekaran et al. 2017) and was carried out with the following hyperparameter, kappa = 10 (the weight of the metabolomic constraints). The change of each metabolite by time was fitted into a linear equation. We took the slope normalized by its intercept as the right-handed side of an FBA problem. The Gurobi mathematical solver was used to solve all optimization problems (Gurobi Optimization, LLC 2023). The objective inference approach performs equally well using other metabolic reconstructions, including Recon2 and Recon3D, and is compatible with other methods for single-cell transcriptomics integration such as COMPASS (Wagner et al. 2021) (Figure S12).

### Flux balance analysis

Two types of models were used for flux prediction: unconstrained models with 52 single objectives (Table 2) and omics-constrained models without objectives. To generate flux predictions with the two models, we applied parsimonious flux balance analysis (pFBA) assuming the minimization of total costs of metabolic enzymes (Jenior et al. 2020). Since metabolites can be in multiple compartments in GEMs, we created demand reactions which are the total amount of those across various compartments, for example, total ATP equals the sum of ATP in cytosol, nucleus, and mitochondria.

### Meta-learner regressor models

A meta-learner model is formed by a set of several candidate models. In this work, we constructed meta-learner models by a linear combination of four different regressors. We chose candidate models including ElasticNet, Lars, Lasso, and stepwise linear regression. To choose hyper-parameters for the first three regressors, five-fold cross-validation was employed to find the best hyper-parameters. Next, we trained the models with five-fold cross-validation and recorded the predictions estimated by the candidate models. Lastly, a linear regression model relied on the predictions from the four regressors as inputs to fit the outcome and estimated coefficients. The coefficients were eventually used to weigh the coefficients of metabolic objectives inferred by the four regressors. A key rationale for choosing a meta-learner regressor model comprising four different linear models over other machine/statistical learning methods was to obtain non-negative coefficients for metabolic objectives. Further meta-learner regression is known to be more numerically stable and helps summarize metabolic objectives from the four individual models (van der Laan, Polley, and Hubbard 2007). The performance of different regression models was evaluated based on correlations between constrained models and fluxes predicted with the inferred coefficients of metabolic objectives. Pearson correlations were 0.755 in the single-cell embryogenesis study, 0.489 in the single-cell cell-cycle study, and 0.702 in the cell quiescence study by using meta-learner regressors (p-value < 10^-16^).

### Analysis of cell-specific metabolic objectives

#### Significant metabolites

We applied a non-parametric method, Mann–Whitney U test, from Scipy to compare metabolic objectives among cell types. Significant metabolites with p-values lower than 0.05 were selected for visualization. Significant metabolites with fold changes higher or lower than 1 were marked with colors corresponding to each cell state/type on the x-axis.

#### Objective entropy

The entropy of metabolic objectives was quantified with the equation *H* =− Σ*_k_* (*p_k_ log*(*p_k_*)) in which *p_k_* was the inferred coefficient of metabolite *k* divided by the total count of metabolites of interest. In the comparisons of entropy between groups, we relied on one-way ANOVA tests to compute p-values and identify significance with 0.05 as a threshold.

#### Distances to the biomass objective

Euclidean distances from inferred objectives to the Recon1’s biomass objective were calculated based on coefficients. Once the biomass precursor was selected by inferred metabolic objectives, we broke down the equation and multiplied each coefficient of the biomass components by the coefficient assigned to the biomass precursor. These values were then reintegrated with other selected metabolite coefficients. Subsequently, the distances of different cell types/states were statistically compared with the T-test.

#### Metabolic trade-offs, Pareto optimality, and traits

Pareto optimality was constructed by flux values of candidate objectives which were transformed into proportions. The flux solutions were generated with a uniform sampling of 10,000 combinations of coefficients between 0 and 1 of metabolites selected by at least 50% of cells/samples. The optimal points were selected by using a Python package, Orthogonal Array package (oapackage) (Eendebak, Pieter Thijs, and Alan Roberto Vazquez. 2019). pFBA took these coefficients to simulate fluxes without constraints. Simultaneously, we converted objective coefficients into proportions and compared them to the Pareto optimality. Next, we integrated the proportions of fluxes of candidate objectives and the proportions of inferred coefficients. PCA then reduced the data dimension and automatically displayed the metabolic traits. The Scree plots showing the variances of each principal component guided us to pick the best number of principal components with the Elbow method. We visually identified the vertices of PCA polygons and extracted their metabolite proportions to represent metabolic traits. In the 3-dimensional Pareto analysis, we applied a Python package, scikit-spatial (Hynes n.d.), to fit the 3-dimensional plane and calculate the distance from each point to the plane. Using uniform random sampling, we constructed the distance from the random coefficients to the 3-dimensional planes as a control. T-test was then used to quantify significance.

#### Metabolic tasks of each metabolic traits

The allocations of the metabolites in metabolic traits were employed to build flux allocations. For example, glycine and NMN were allocated 0.3 and 0.7, respectively, in a trait. The flux allocation of the trait was the summation of glycine-optimized flux solution multiplied by 0.3 and NMN-optimized flux solution multiplied by 0.7 which were eventually converted into absolute values. To account for the allocation of each subsystem, we normalized the linear combination of flux solutions by the summation of all the fluxes. We next generated 1000 random allocations of metabolites shown in traits and built the linear combination of flux allocations as controls. By comparing to the control flux allocations, we labeled reactions with flux allocations higher than the upper bound of 95% confidence interval. Relying on the method, we calculated task scores with the number of labeled reactions in each subsystem normalized the total number of reactions in the subsystem. The subsystems identified as metabolic tasks in each trait shown in Table S1 had task scores higher than mean of task scores of all traits.

## Acknowledgments

National Institutes of Health grant R35 GM13779501 (SC) Camille and Henry Dreyfus Foundation (SC)

## Data availability

All datasets are provided in figures, supplementary tables and in the Github page

## Author contributions

Conceptualization: SC

Investigation: DL

Funding acquisition: SC, JZ

Project administration: SC, JZ

Supervision: SC, LZ, JZ

Writing – original draft: SC, DL

Writing – review & editing: SC, LZ, JZ, DL

## Supplementary Materials

### Data sources

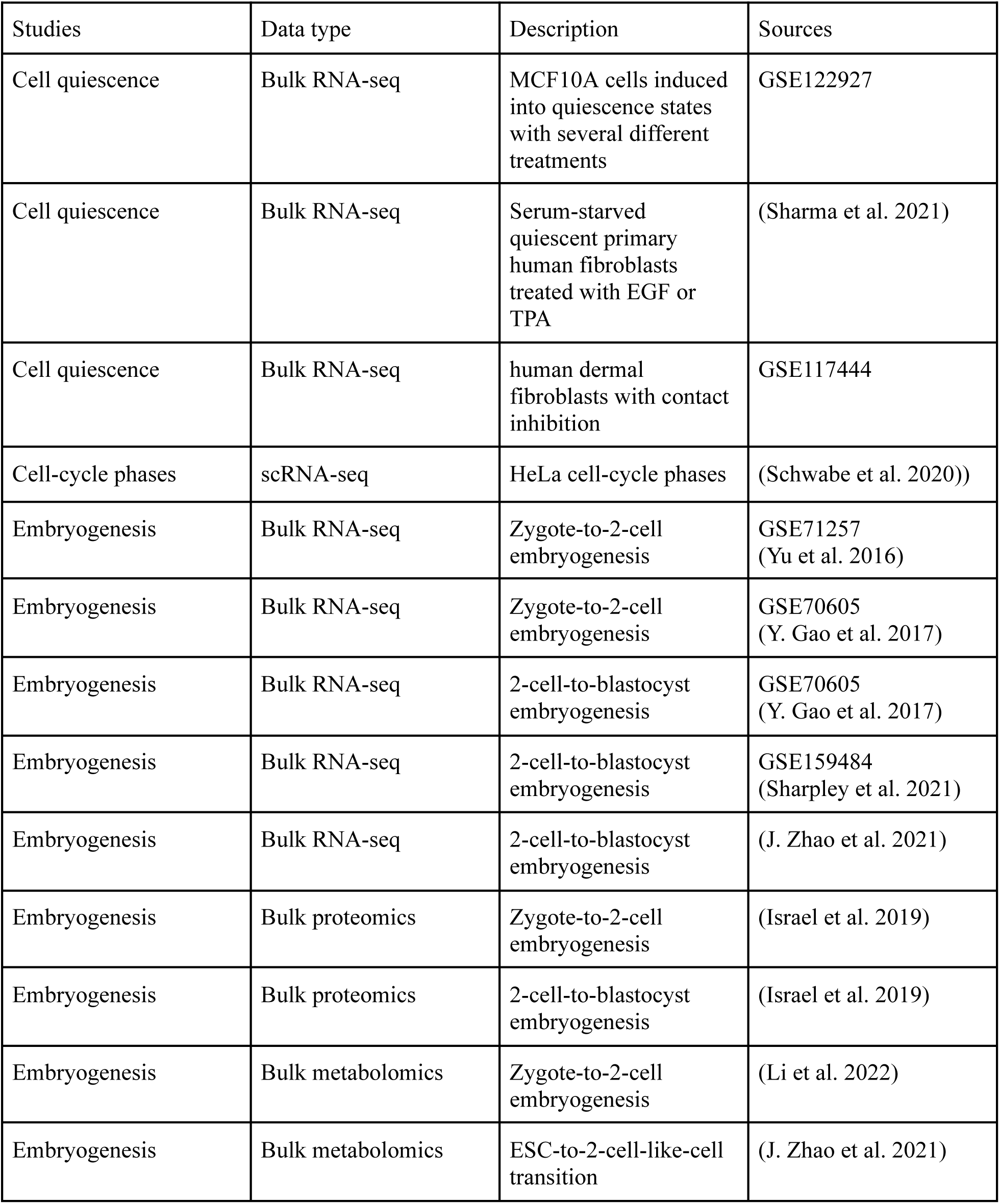
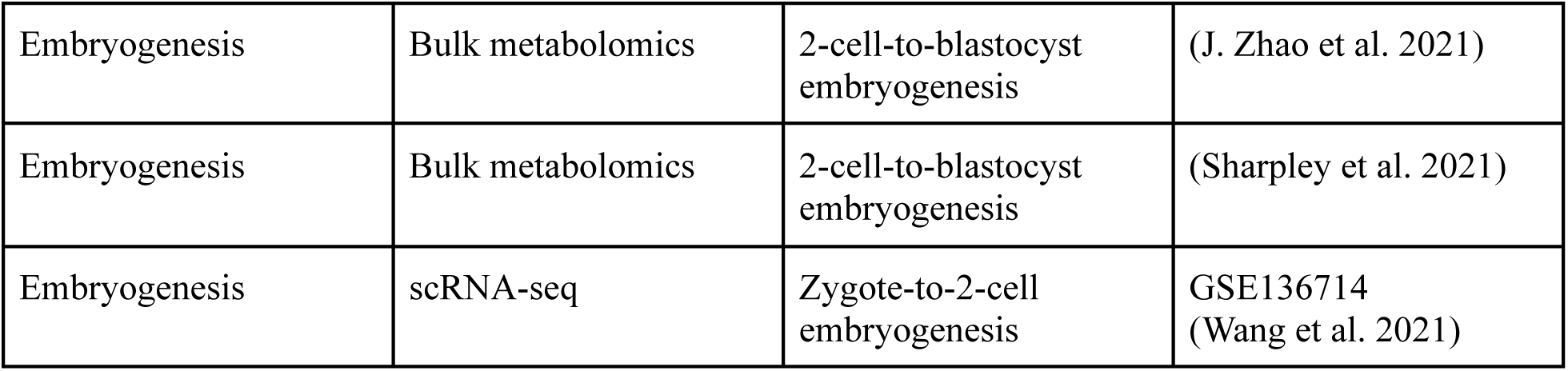

**Supplementary Figure 1.**
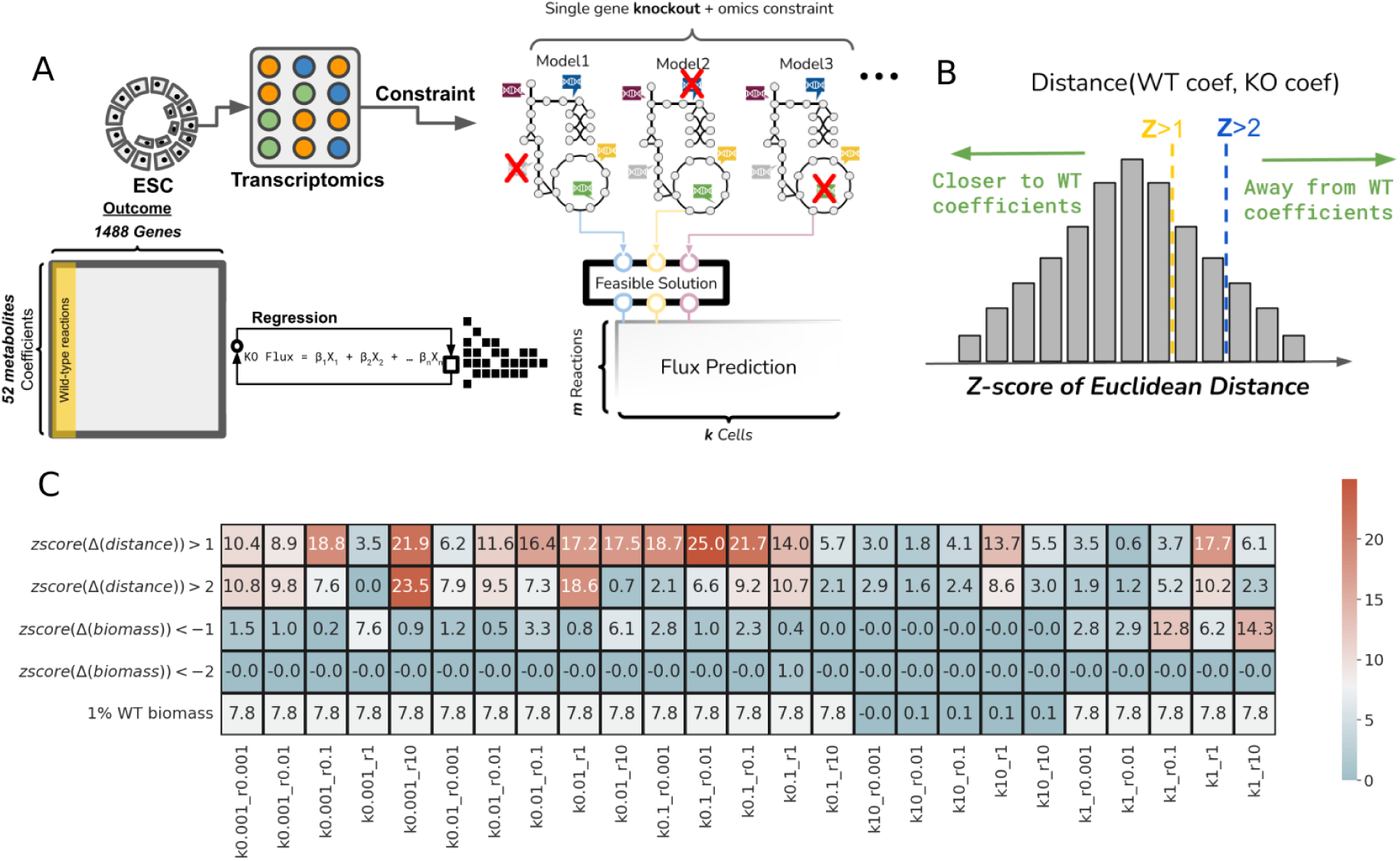
Evaluation of metabolic objectives on gene essentiality. (A) The workflow of using metabolic objectives to predict gene essentiality starts from inferring transcriptomics-based metabolic objectives of embryonic stem cells (ESC) (Y. Zhang et al. 2013). FDR p-values less than 0.05 was used to select essential genes from genome-wide CRISPR screens collected in Shohat et al. and Tzelepis et al. (Shohat and Shifman 2019; Tzelepis et al. 2016). Wild-type (WT) flux prediction was constrained with the ESC transcriptomics without optimizing production of any metabolites. Likewise, we applied the gene-protein-reaction rule to remove single gene in each knockout (KO) model. Subsequently, we inferred the metabolic objectives of the WT and KO fluxes. (B) We compared wild-type to knockout objective coefficients with Euclidean distances. The essential genes were identified using the Z-scores of the distances with threshold 1 or 2. (C) The method mentioned in (A) were compared to traditional methods. The last 3 rows in the heatmap were calculated with the biomass objective values estimated with a model constrained with the ESC transcriptomics which optimized the biomass objective. Essential genes were identified with the differences of the biomass objective values between KO and WT models. The *zscore*(Δ*biomass*) <− 1 and the *zscore*(Δ*biomass*) <− 2 considered genes essential with the Z-scores of the differences less than -1 or -2 as thresholds. In contrast, single gene knockouts resulting in the biomass objective values less than 1% of the value in the WT model was shown in the last row. Hypergeometric test was employed to evaluate the predictions and shown in the heatmap with negative Log-10 p-values.

**Supplementary Figure 2.**
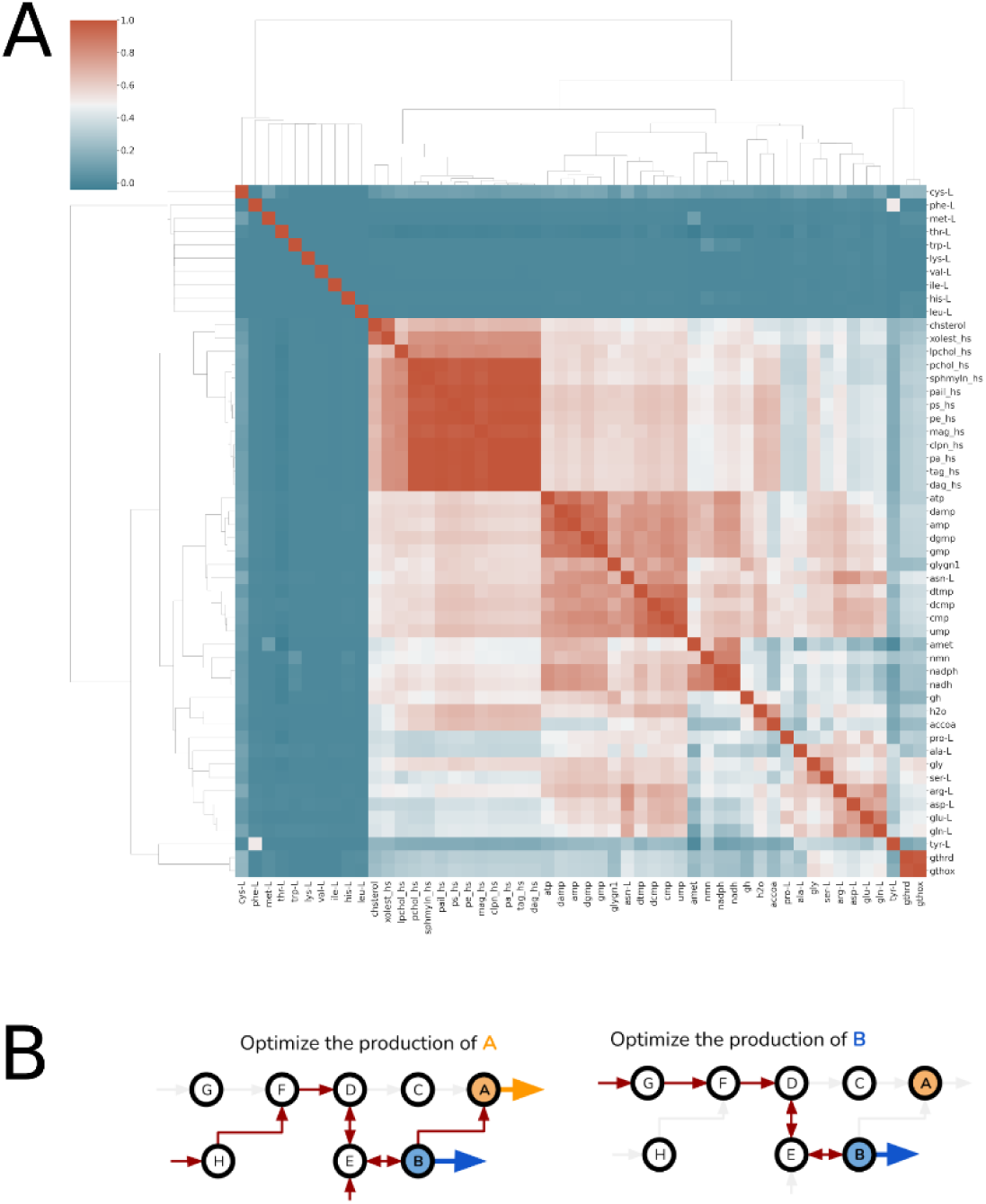
Optimal flux solutions optimizing individual metabolites. (A) Pearson correlation of flux allocation optimizing single metabolite clustered lipids, nucleotides, growth-boosting amino acids, and glutathione. However, amino acid including cysteine, leucine, and methionine were dissimilar and different from other metabolites. (B) An illustration of unconstrained models optimizing different metabolites sharing similar flux solutions.

**Supplementary Figure 3.**
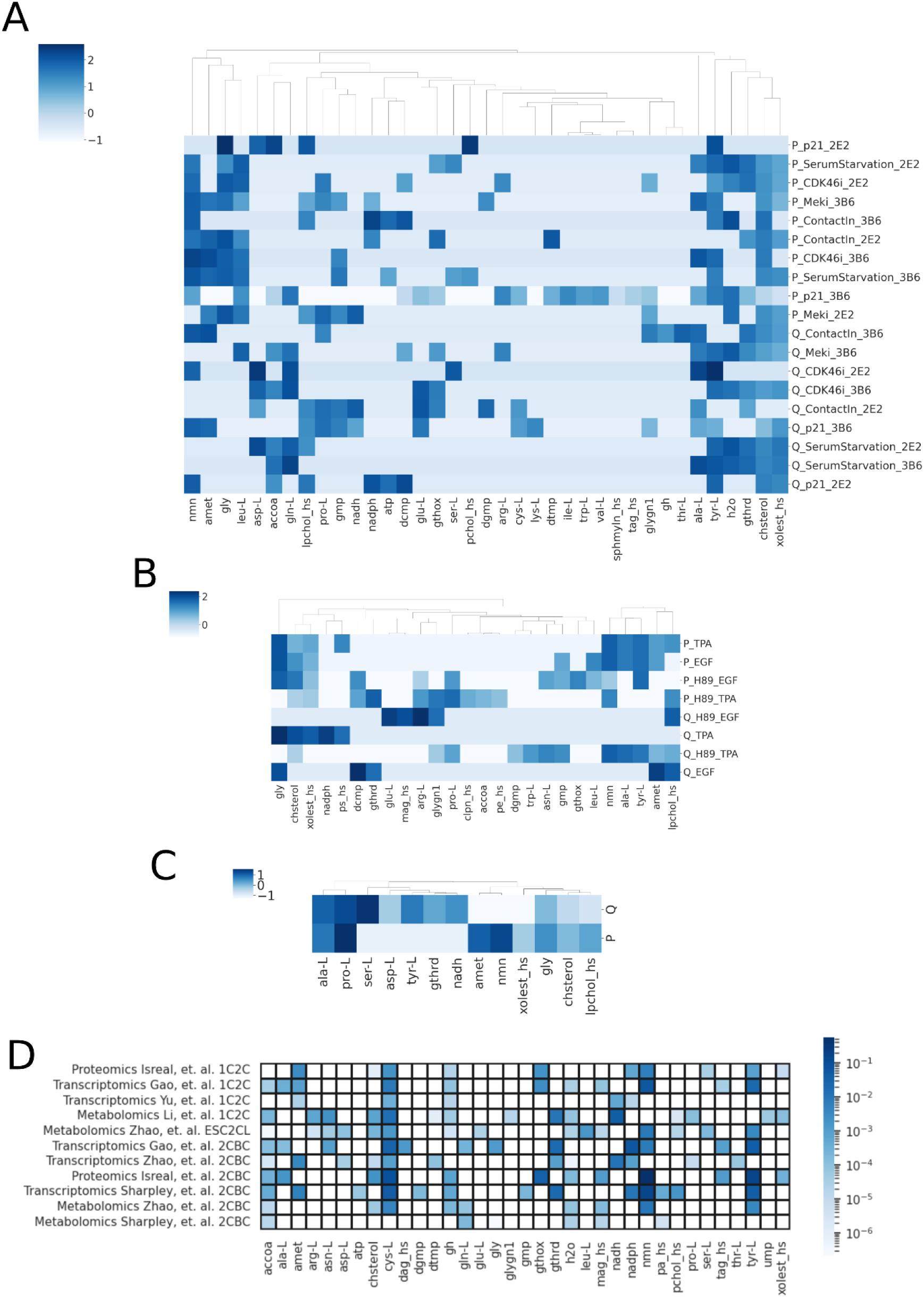
Clustering of proliferative and quiescent states. (A-C) The inferred metabolic objectives shown in Figure 2B and D predicted in quiescent (shown with Q_ prefix) versus proliferative (shown with P_ prefix) states from (A) MCF10A epithelial cells, (B) primary human fibroblasts, and (C) dermal fibroblasts (Min and Spencer 2019; Mitra et al. 2018; Sharma et al. 2021). (A). The four different conditions compared sorted cells with high (quiescent, Q) or low p21 (proliferative, P) expression, with (Q) or without (P) MEK inhibition (Meki), with (Q) or without (P) serum starvation, and with (P) or without (Q) contact inhibition (Min and Spencer 2019). (B) Quiescent primary human fibroblasts under serum starvation were exposed to epidermal growth factor (EGF) or the phorbol ester (12-O-tetradecanoylphorbol-13-acetate, TPA), with or without the MSK/PKA inhibitor H89 to induce proliferation (Sharma et al. 2021). (C) Dermal fibroblasts were treated with contact inhibition (Q) or not (P) for seven days (Mitra et al. 2018). (D) Metabolic objectives were inferred with the bulk-omics datasets of the 1C2C and the 2CBC transitions. In Figure 4B, metabolites having non-zero coefficients in at least two genomics-based objectives and at least one metabolomics dataset were shown in the heatmap. This figure shows the completed version of Figure 4B displaying metabolic objectives inferred with the bulk-omics datasets of the 1C2C and the 2CBC transitions.

**Supplementary Figure 4.**
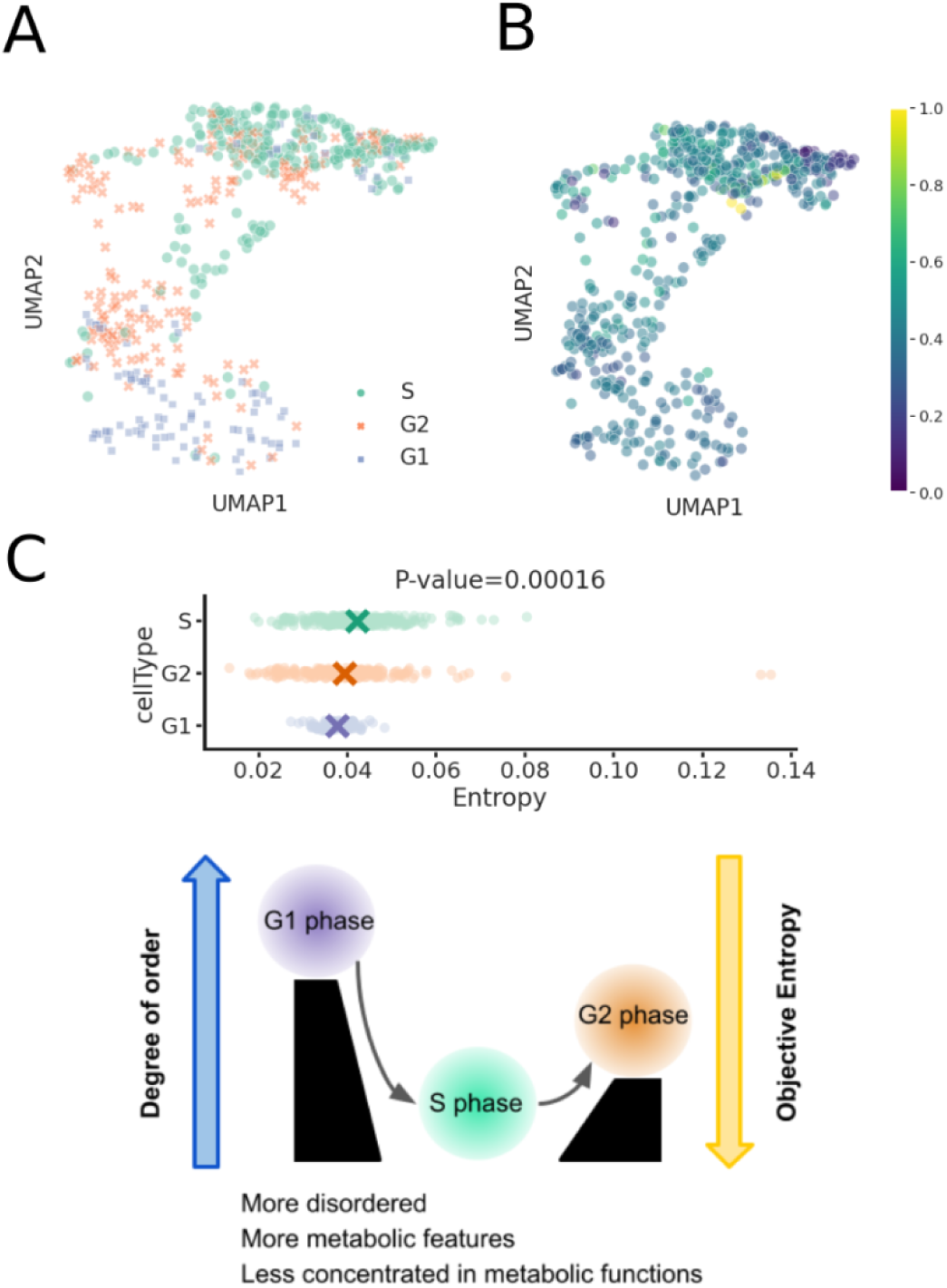
Dimension-reduction of cell-cycle metabolic objectives and the objective entropy. (A) UMAP of cell-cycle metabolic objectives separated by cell-cycle phases. The low-dimensional projection placed the S, G2, and G1 phases from the top to the bottom of the plot. (B) The low-dimensional projection of metabolic objectives were overlaid with objective entropy which displayed an increase trend from the bottom to the top followed by a sudden drop. (C) One-way ANOVA test suggested significant different entropy in the three cell-cycle phases. The S phase had the greatest entropy followed by the G2 phase and then the G1 phase. (D) The changes of objective entropy can be explained by the allocations of metabolic functions in different cell-cycle phases. The highest entropy in the S phase indicated the multitasking nature of its metabolic network.

**Supplementary Figure 5.**
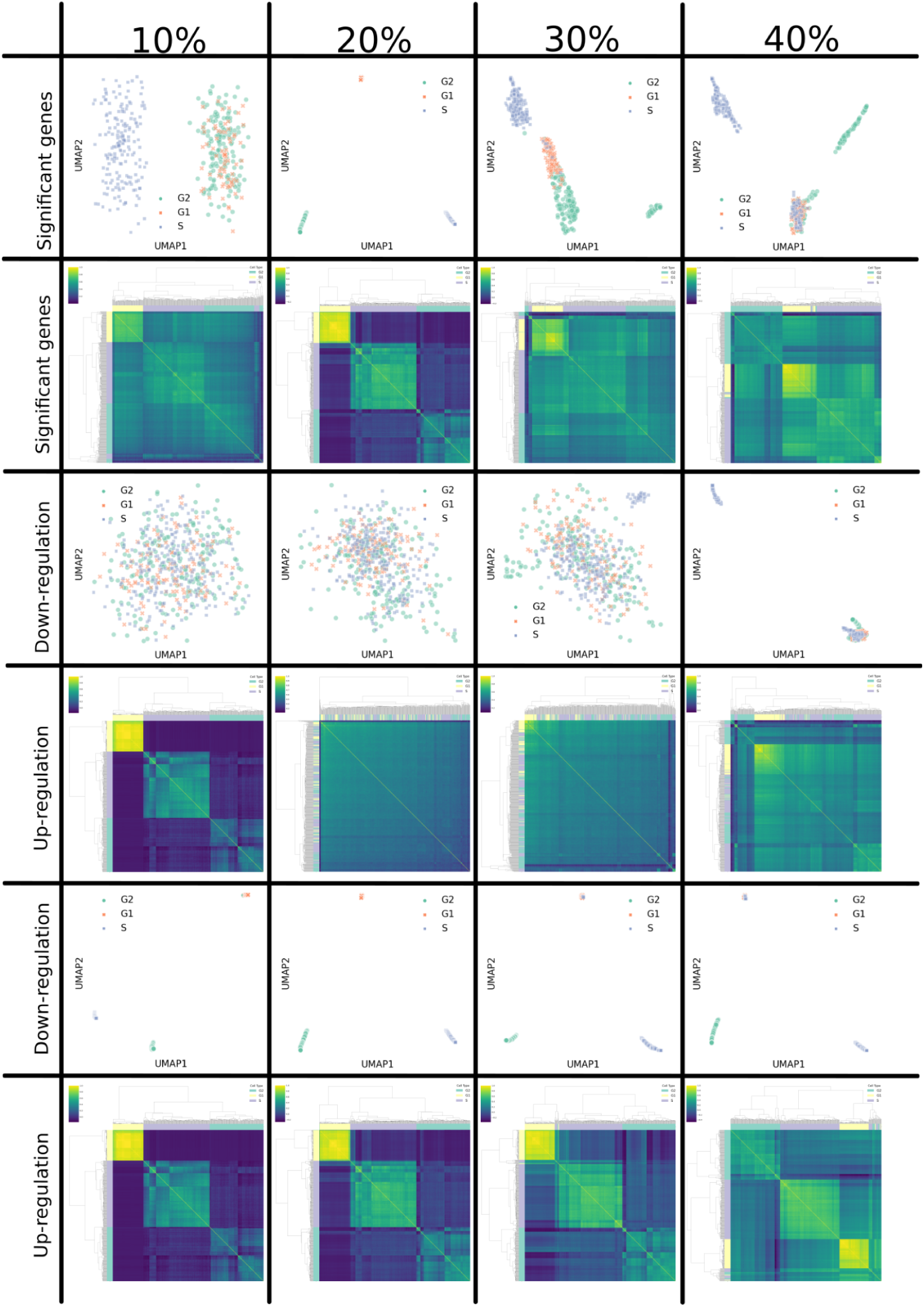
Threshold selections for up- and down-regulated metabolic genes in single-cell cell-cycle data. (Top two rows) The first two rows are UMAP and hierarchical clustering of Pearson correlation based on the classification of metabolic genes. (The rest four rows) Up- and down-regulated metabolic genes were individually shown with UMAP and hierarchical clustermap. Only using 40% as a threshold could separate the S phase in up-regulated genes.

**Supplementary Figure 6.**
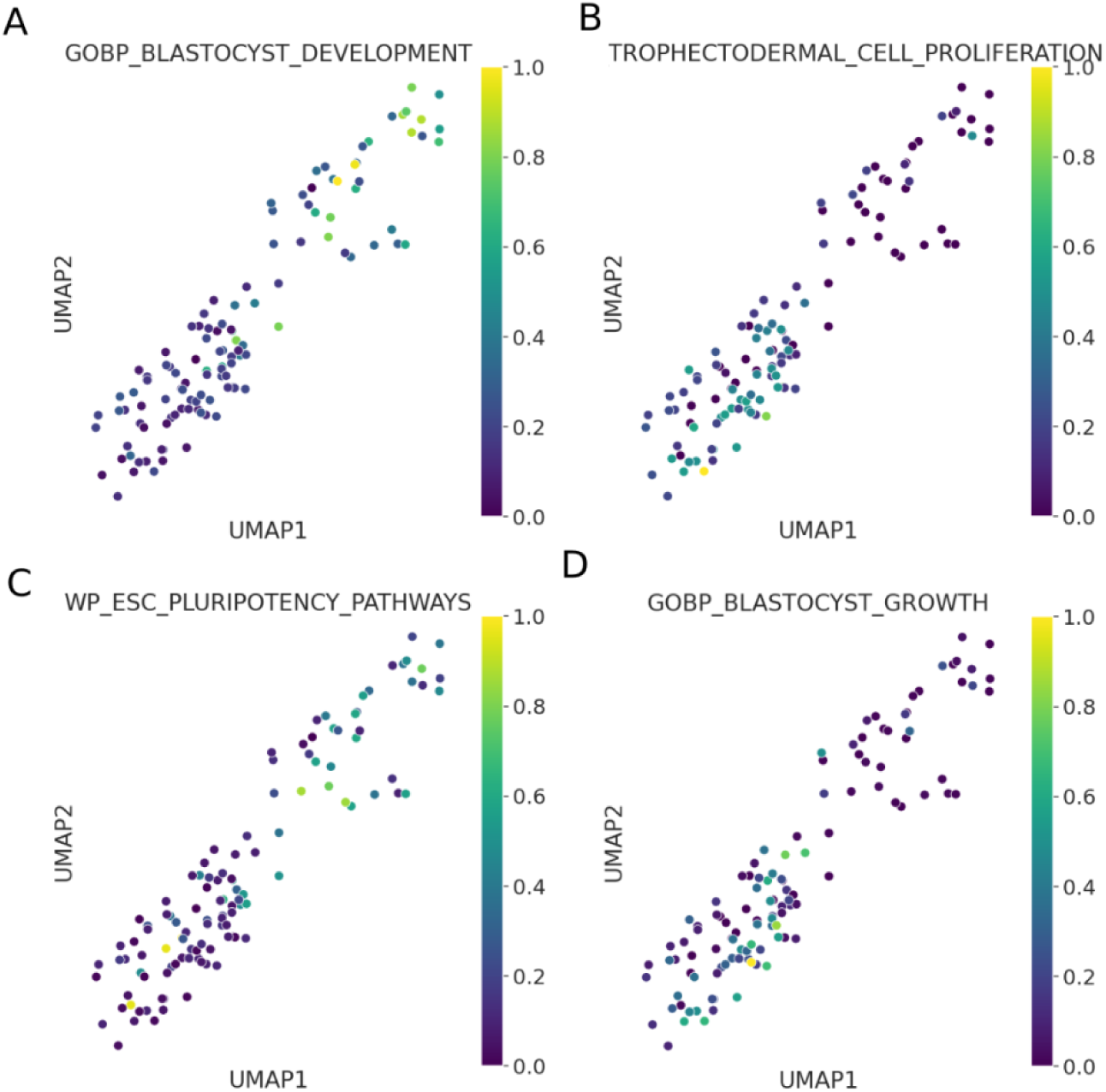
Overlaying enrichment analysis over UMAP of metabolic objectives. We applied hypergeometric test to 4 sets of gene sets from GSEA MsigDB database including (A) blastocyst development, (B) trophectoderm cell proliferation, (C) ESC pluriotency pathways, and (D) blastocyst growth. Negative log p-values of the hypergeometric test were further linearly scaled in the range from 0 to 1 for visualization using the Python package matplotlib,normalize. Higher values imply higher enrichment.

**Supplementary Figure 7.**
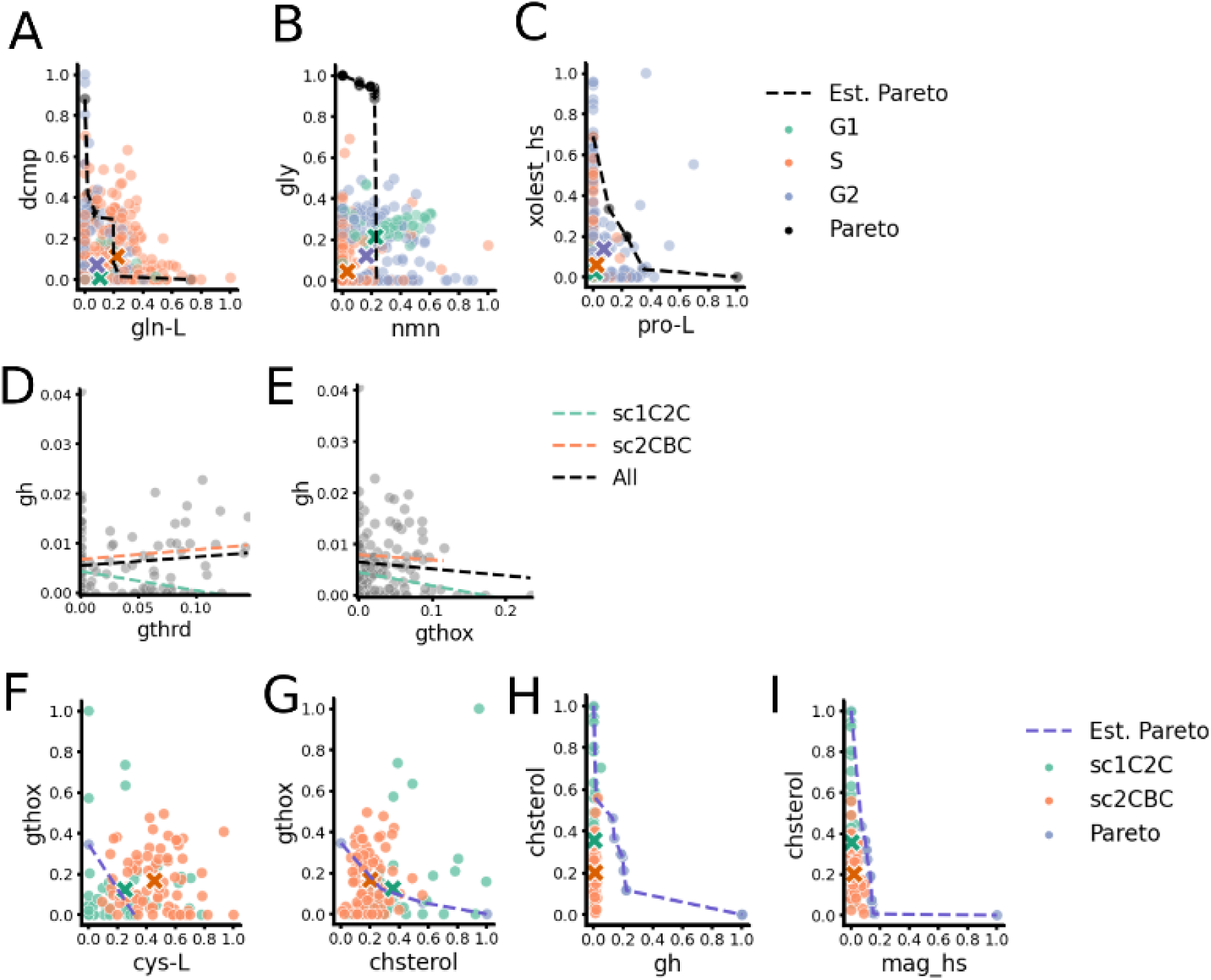
Metabolic trade-offs in the cell cycle and early embryogenesis. (A-C) The Pareto analysis of significant metabolites in each cell-cycle phase. (A) The S phase was the closest compared to the G1 and G2 phases to the Pareto front of dCMP and glutamine which were significantly higher in S phase. Likewise, the G1 phase and the G2 phase were closest to (B) the Pareto fronts of glycine and NMN and the (C) Pareto front of cholesterol ester and proline, respectively. (D, E) The relationship of coefficients between the biomass objective (gh) and reduced or oxidized glutathione in the 1C2C and 2CBC transitions. (F-I) The 1C2C and 2CBC transitions were both closer to the Pareto fronts of four pairs of metabolites including (F) oxidized glutathione against cysteine, (G) oxidized glutathione against cholesterol, (H) cholesterol against the biomass precursor, and (I) cholesterol against monoacylglycerol 2.

**Supplementary Figure 8.**
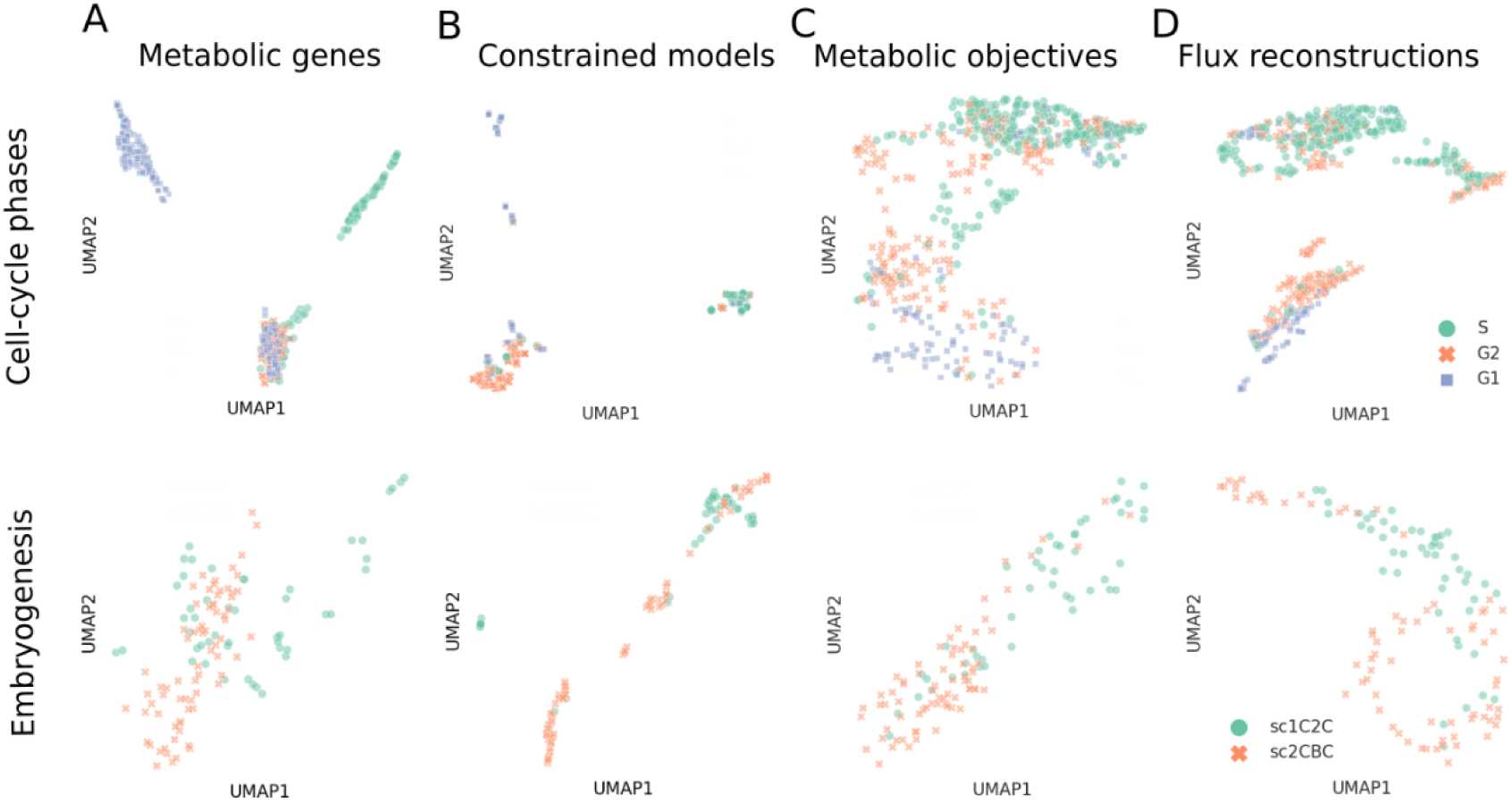
UMAP visualization of metabolic objectives and reconstructed fluxes. UMAP plot of (A) up- and down-regulated metabolic genes, (B) constrained models without objectives, (C) inferred metabolic objectives, and (D) reaction fluxes predicted with the regression models. The top row is cell-cycle phases and the bottom is embryogenesis. This shows that the inferred objectives can better summarize and visualize the metabolic differences between cell types than the input transcriptomics data.

**Supplementary Figure 9.**
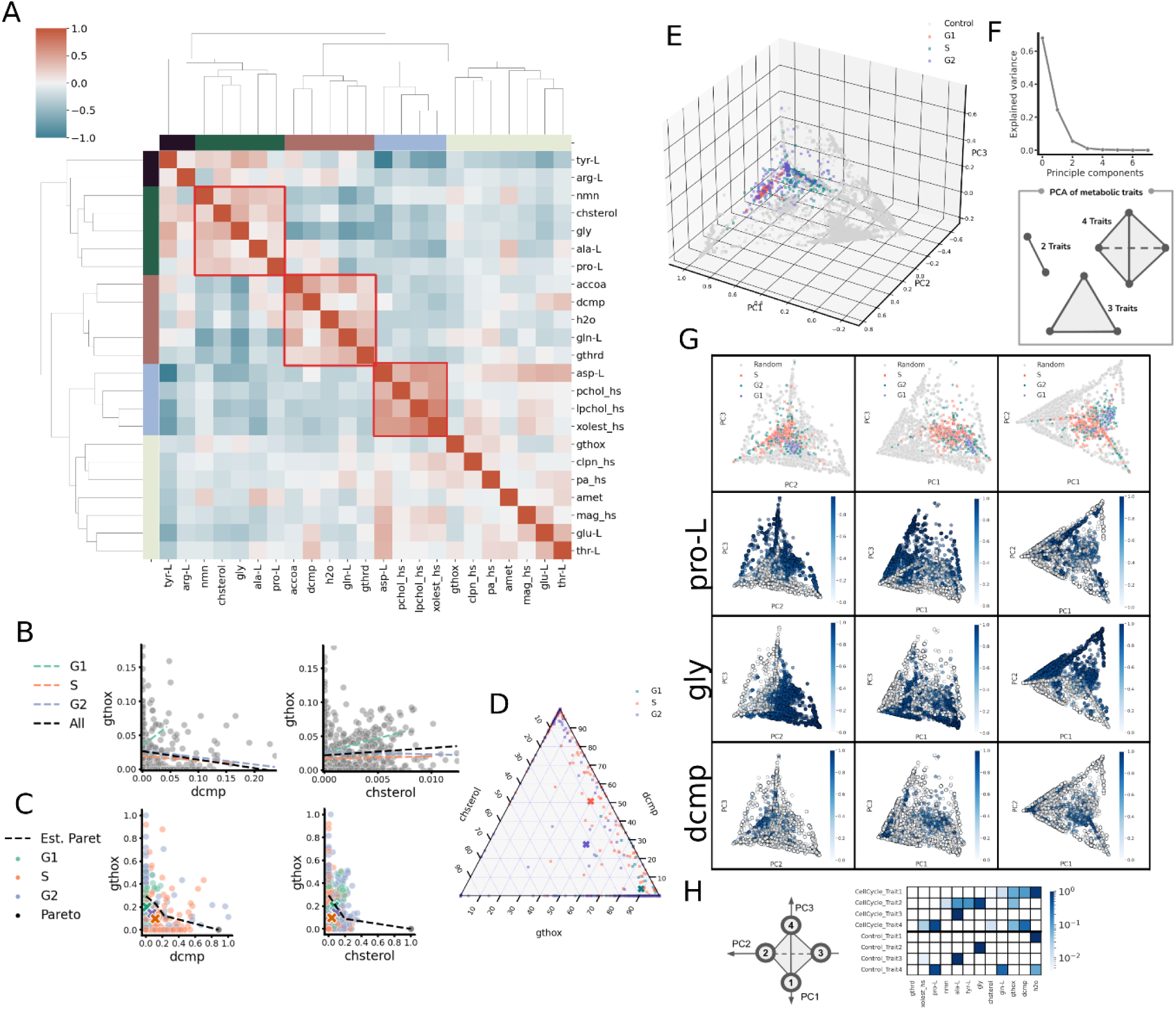
Metabolic objective trade-offs in cell-cycle phases. (A) Normalized Pearson correlations of objective coefficients showed the relationships among candidate metabolites. Negative correlations shown in blue implied potential trade-offs. (B) The relationships between reduced glutathione and oxidized glutathione, cholesterol, and dCMP were shown in the scatter plots and fitted by each cell-cycle phase. (C) Normalized objective coefficients show Pareto optimalities. Objectives estimated by simulating fluxes with random coefficients are shown in black. The mean coefficients of each cell-cycle phase are shown as “X”. A closer mean coefficient to the pareto front implies that the traits are simultaneously optimized and form a trade-off. (D) The proportion of coefficients among cholesterol, dCMP, and the summation of oxidized and reduced glutathione was shown with a ternary plot. Each axis represents the percentage of a metabolite coefficient allocated in single cells. (E-H-J) The top 50% of candidate metabolites were selected to model the ideal objective fluxes using random objective coefficients. Subsequently, the flux values were normalized to proportions that quantify the distribution of metabolite allocations within single cells. (E) A PCA plot automatically extracted 4 traits, according to (F) the Elbow method, by integrating the normalized ideal objective fluxes with the normalized inferred coefficients. (G) Projections of the 3-dimensional PCA plot are shown in the top row. In the following rows, the proportions of dCMP, glycine, alanine, and proline are overlaid on the PCA projections, respectively. (J) The single cells (CellCycle_Trait) or control data points (Control_Trait) closest to the theoretical traits were selected to show their proportions with a heatmap. The suffix numbers correspond to the numbers labeled in the left illustration. These findings suggest that distinct metabolic function allocations have been optimized for the G1, G2, and S phases. These patterns highlight the potential trade-offs among redox, nucleotide, and lipid metabolism, shaping the metabolic characteristics of cell-cycle phases.

**Supplementary Figure 10.**
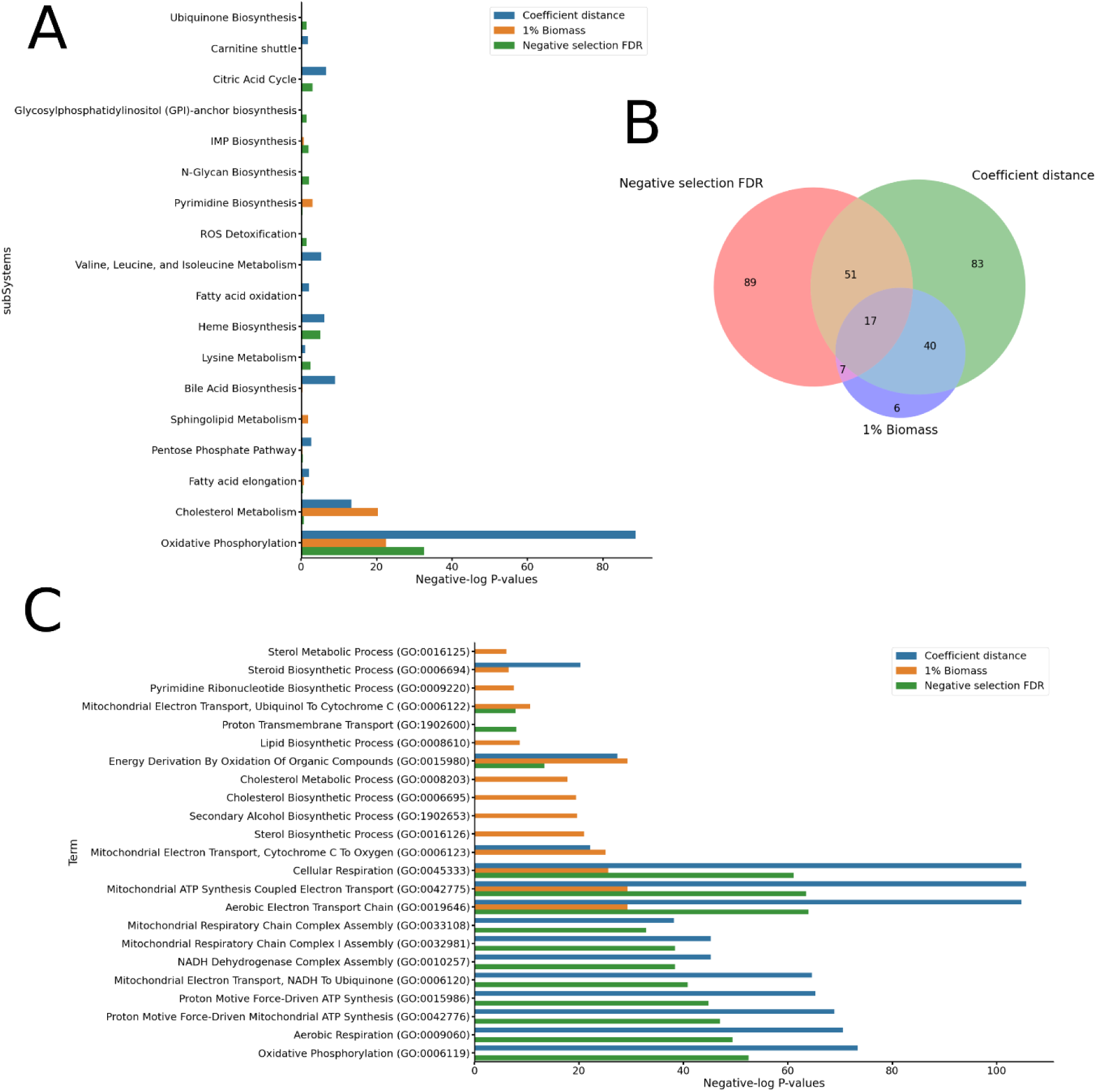
Enrichment analysis of essential genes in ESC. Comparison of the essential genes identified experimentally with those obtained using traditional biomass objective and the objective inference method. Genes that reduce the biomass objective values by less than 1% of the wild type value and those genes with Z-score of coefficient distances from WT model to KO model greater than 1 were selected respectively for enrichment analysis. Enrichment using (A) subsystems of GEM and (C) gene ontology (GO) terms are shown. The overlap of the gene lists from the two methods with the experimental data (negative selection of the CRISPR-Cas9 screen) are shown in the Venn diagram (B). The objective inference method correctly identified genes in heme synthesis, oxidative phosphorylation and TCA cycle which were also seen experimentally but were missed by the traditional biomass objective.

**Supplementary Figure 11.**
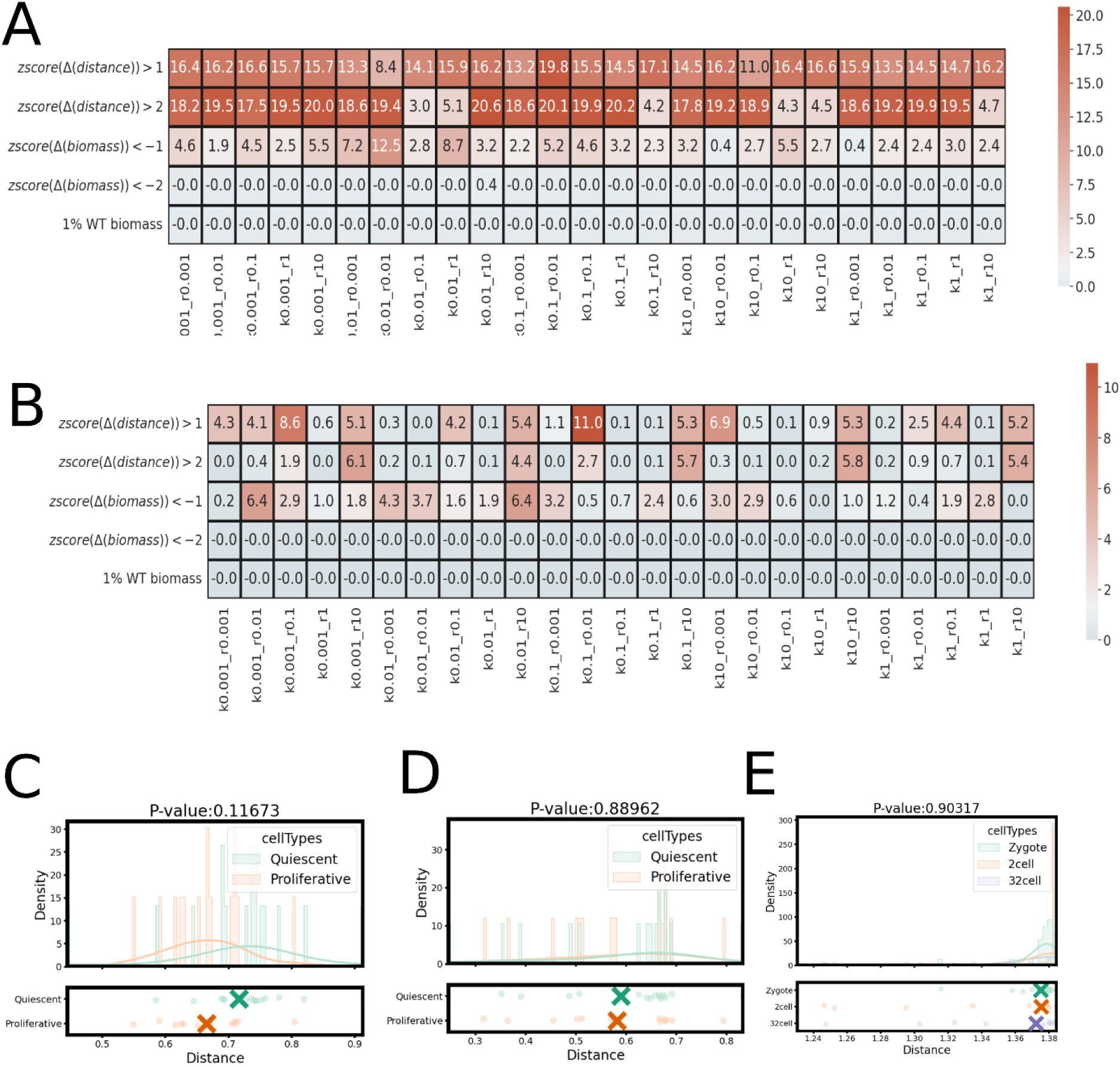
Inference of metabolic objectives with different model settings. Metabolic objectives inferred with (A) Recon 2.2 and (B) Recon 3D were evaluated with the CRISPR-Cas9 screen in ESC. The method follows the workflow in Figure S1A. In the first two rows, we compared wild-type to knockout objective coefficients with Euclidean distances. The essential genes were identified using the Z-scores of the distances with threshold 1 or 2. The last 3 rows in the heatmap were calculated with the traditional biomass objective values estimated with a model constrained with the ESC transcriptomics which optimized the default biomass objective used in (A) Recon 2.2 and (B) Recon 3D. Essential genes were identified with the differences of the biomass objective values between KO and WT models. The Δ*biomass* < 0 considered genes essential with the difference greater than 0. Essential genes were identified with the differences of the biomass objective values between KO and WT models. The Δ*biomass* < 0 considered genes essential with the difference greater than 0. The *zscore*(Δ*biomass*) <− 1 and the *zscore*(Δ*biomass*) <− 2 considered genes essential with the Z-scores of the differences less than -1 or -2 as thresholds. In contrast, single gene knockouts resulting in the biomass objective values less than 1% of the value in the WT model was shown in the last row. FDR p-values less than 0.05 was used to select essential genes from genome-wide CRISPR screens collected in Shohat et al. and Tzelepis et al. (Shohat and Shifman 2019; Tzelepis et al. 2016). Hypergeometric test was employed to evaluate the predictions and shown in the heatmap with negative Log-10 p-values. (C-D) The Euclidean distance from the inferred metabolic objective to the default biomass objectives used in (C) Recon 2.2 and (D) Recon 3D confirmed that the proliferative states were closer to the biomass objectives. (E) We applied COMPASS to constrain Recon 1 with the single-cell transcriptomics of embryogenesis. The constrained models were then followed the same workflow mentioned in Figure 1 to infer metabolic objectives. As a result, the Euclidean distance from the inferred metabolic objective to the Recon 1’s biomass objectives confirmed that the 32-cell (blastocyst) were closer to the biomass objectives, although the trend was not statistically significant.

**Table S1.** Metabolic functions corresponding to each embryo trait.

| Embryo Trait 1 | Embryo Trait 2 | Embryo Trait 3 |
| --- | --- | --- |
| Pyrimidine Biosynthesis | D-alanine metabolism | beta-Alanine metabolism |
| Methionine Metabolism | Salvage Pathway | Arginine and Proline Metabolism |
| Inositol Phosphate Metabolism | R Group Synthesis | Glyoxylate and Dicarboxylate Metabolism |
| Glutathione Metabolism | Fatty acid elongation | Pyruvate Metabolism |
| Glycerophospholipid Metabolism | Tyrosine metabolism | Propanoate Metabolism |
| Triacylglycerol Synthesis | Alanine and Aspartate Metabolism | NAD Metabolism |
| IMP Biosynthesis |  | Urea cycle/amino group metabolism |
| Starch and Sucrose Metabolism |  | Methionine Metabolism |
| Galactose metabolism |  | Tryptophan metabolism |
| Sphingolipid Metabolism |  | Folate Metabolism |

